# Alpha Oscillatory Activity Causally Linked to Working Memory Retention: Insights from Online Phase-corrected Closed-loop Transcranial Alternating Current Stimulation (tACS)

**DOI:** 10.1101/2021.05.23.445322

**Authors:** Xueli Chen, Ru Ma, Wei Zhang, Qianying Wu, Ajiguli Yimiti, Xinzhao Xia, Jiangtian Cui, Ginger Qinghong Zeng, Junjie Bu, Qi Chen, Nancy Xiaonan Yu, Shouyan Wang, Zhi-De Deng, Alexander T. Sack, Myles Mc Laughlin, Xiaochu Zhang

## Abstract

Although previous studies have reported correlations between alpha oscillations and the “retention” sub-process of working memory (WM), causal evidence has been limited in human neuroscience due to the lack of delicate modulation of human brain. Conventional tACS is not suitable for demonstrating the causal evidence for parietal alpha oscillations in WM retention because of its inability to modulate brain oscillations within a short period (i.e., the retention sub-process). Here, we developed an online phase-corrected closed-loop transcranial alternating current stimulation (tACS) system capable of precisely correcting for the phase differences between tACS and concurrent endogenous oscillations. This system permits both up- and down-regulation of brain oscillations at the target stimulation frequency within a short stimulation period, and is here applied to empirically demonstrate that parietal alpha oscillations causally relate to WM retention. Our experimental design included both in-phase and anti-phase alpha-tACS applied to 39 participants during the retention sub-processes of a modified Sternberg paradigm. Compared to in-phase alpha-tACS, anti-phase alpha-tACS decreased both WM performance and alpha activity. Moreover, the in-phase tACS-induced changes in WM performance were positively correlated with alpha oscillatory activity. These findings strongly support a causal link between alpha oscillations and WM retention, and illustrate the broad application prospects of phase-corrected tACS.

## 1. Introduction

Working memory (WM) is considered to be foundational for a broad range of cognitive functions (*e.g.*, the capacity for general intelligence, categorization, retrieving selected long-term memories, language learning, etc.) [1]. WM enables the maintenance, manipulation, and retrieval of mental representations, as well as the use of this information in goal-directed behaviors [1]. Because of its essential role in human cognition, investigating the neural mechanisms underlying WM has been a focus of neuroscience research for decades [2, 3].

WM can be subdivided into three fundamental sub-processes: encoding, retention, and retrieval [4]. Neural oscillations in the alpha frequency band (8-13Hz) have been associated with WM retention [5]: alpha activity in the occipitoparietal areas increases during memory retention [6–10]. Previous studies also reported that alpha power increased parametrically with memory load during the retention interval [6, 11]. However, as these previous studies only used MEG or EEG, their inferences are correlational in nature; there have not been direct experimental investigations which successfully demonstrated a causal role for alpha oscillations in WM retention in humans.

Transcranial alternating current stimulation (tACS) may provide us with the opportunity to experimentally investigate a causal role for alpha oscillations in WM retention, owing to its ability to entrain naturally occurring neural oscillations based on externally-applied, sinusoidal electric fields at a targeted frequency [12–15]. It should be noted that results from previous alpha-tACS studies are at the center of a controversial debate in which some studies failed to replicate successful entrainment effects of alpha-tACS [16]. This discrepancy across alpha-tACS studies may be related to the fact that previous efforts have (to our understanding) not accounted for phase differences between an applied sinusoidal waveform and the concurrent, endogenous oscillations occurring in the targeted brain regions [17]. Although direct evidence based on monitoring oscillatory neural activity in humans is absent, recent computational model [18], animal study [19], and indirect experimental evidence correcting for the phase differences between tACS and peripheral tremor in Parkinson’s patients [20] suggest that the phase differences between tACS and concurrent brain oscillations are impactful for determining the efficacy of tACS (also for determining the robustness of effects and replicability).

Moreover, contrary to the retention sub-process, decreased (rather than increased) alpha oscillations during the encoding and retrieval sub-processes were reported to be beneficial for WM performance [21, 22]. Thus, tACS investigating the causal role for alpha oscillations in WM retention should be time-locked to the specific WM sub-process of interest (i.e., retention but not encoding or retrieval), to avoid the offset effects of conventional tACS applied during the other two sub-processes given that conventional tACS was usually applied continuously for 20-30 minutes.

Given this background, there are two major reasons that conventional tACS methods do not support experimental investigations about the function(s) of alpha oscillations in WM retention: conventional tACS methods cannot correct for phase differences between tACS and endogenous brain oscillations in real-time (“online”), and conventional tACS methods do not support tACS stimulation that can be time-locked to the specific short-time sub-process of WM. These ideas motivated our desire for a tACS technology with the following capabilities: 1) a capacity for online monitoring of endogenous alpha oscillatory activity, to support real-time phase alignment between externally applied tACS and concurrent endogenous alpha oscillations, and 2) a capacity to induce short-duration (within seconds) tACS in a time-locked manner which can be matched to the short-duration retention interval in each trial.

The present study successfully developed a trial-by-trial closed-loop tACS-EEG design that is capable of correcting for the phase differences between applied alpha-tACS and concurrent endogenous oscillations (**Fig. 1** **& 2**), specifically during the retention intervals of a Sternberg WM task [23]. Importantly, this technique allowed us to directly investigate whether alpha oscillations exert any causal impacts on WM retention. Our experimental investigations included alpha-tACS with three different phase differences to the online monitored endogenous alpha oscillations (in-phase tACS, anti-phase tACS, and random-phase tACS, **Fig. 3C**) applied specifically during the WM retention intervals, as well as a control theta-tACS experiment to exclude any general effects of tACS (i.e., independent of frequency) [24]. Fascinatingly, we found that compared to in-phase alpha-tACS, anti-phase alpha-tACS suppressed WM performance, parietal alpha power, and frontoparietal alpha synchronization, and noted that changes in WM performance induced by in-phase alpha-tACS were positively correlated with changes in endogenous alpha oscillatory activity. Ultimately, beyond experimentally demonstrating a causal role for parietal alpha oscillations in WM retention, our results clearly illustrate how phase differences between tACS and concurrent endogenous brain oscillations determine the efficacy and replicability of tACS effects.

**Fig. 1.**
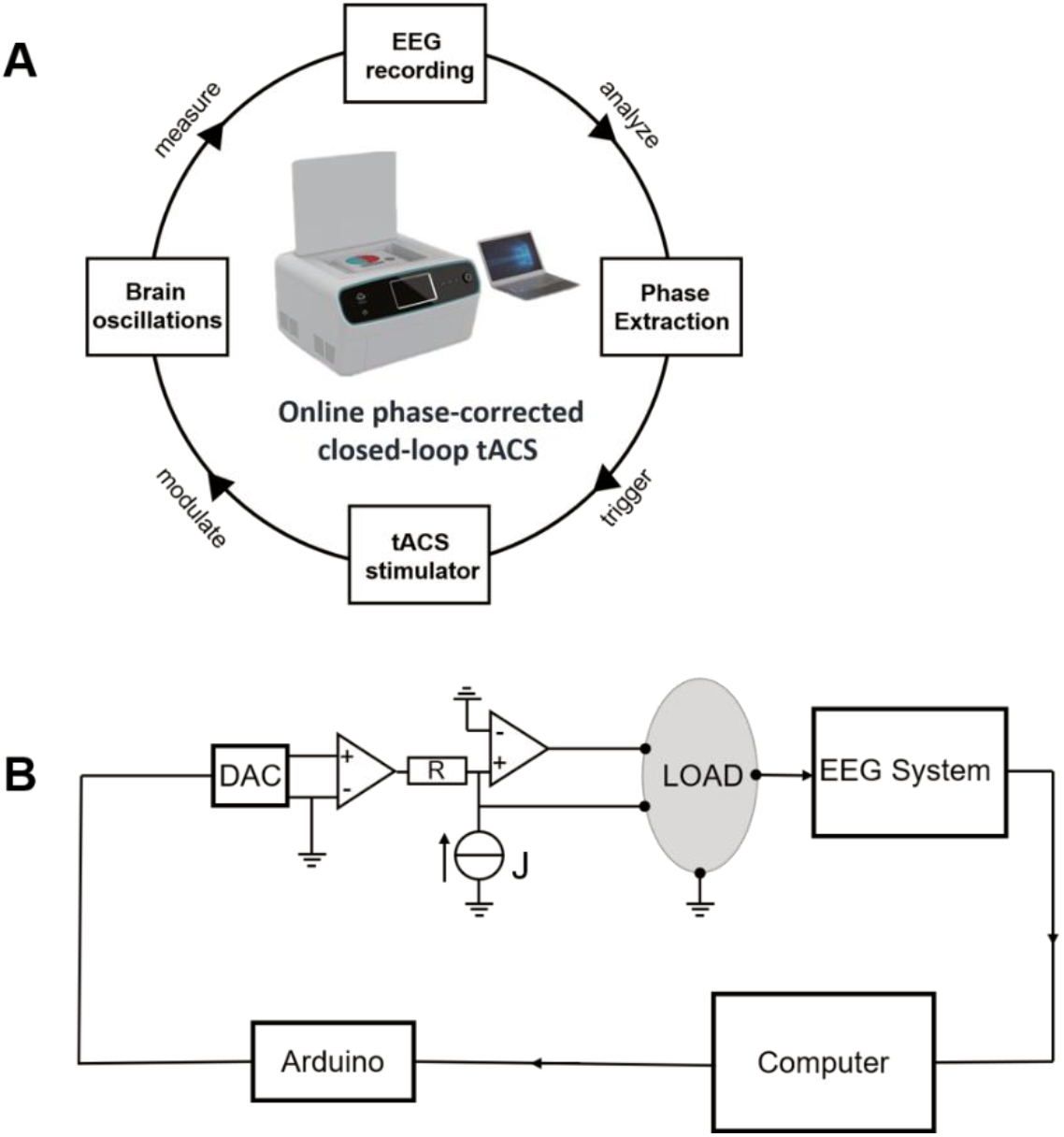
The components of the online phase-corrected closed-loop tACS system and the design of the tACS stimulator. (**A**) The online phase-corrected closed-loop tACS system consists of an EEG instrument measuring brain oscillations, a computer extracting the online phases of brain oscillations to decide the timing of tACS stimulator, and a custom designed tACS stimulator which communicates with the computer to regulate the application of tACS to the human brain. (**B**) The design of the tACS stimulator. The major components of the tACS stimulator include an Arduino Uno microcontroller board, a digital-to-analog converter (DAC), a constant-voltage source (J), and two operational amplifiers. The timing of next tACS (calculated from the computer) is communicated to the Arduino board. The triangle represents the operational amplifier; R represents the resistance.

**Fig. 2.**
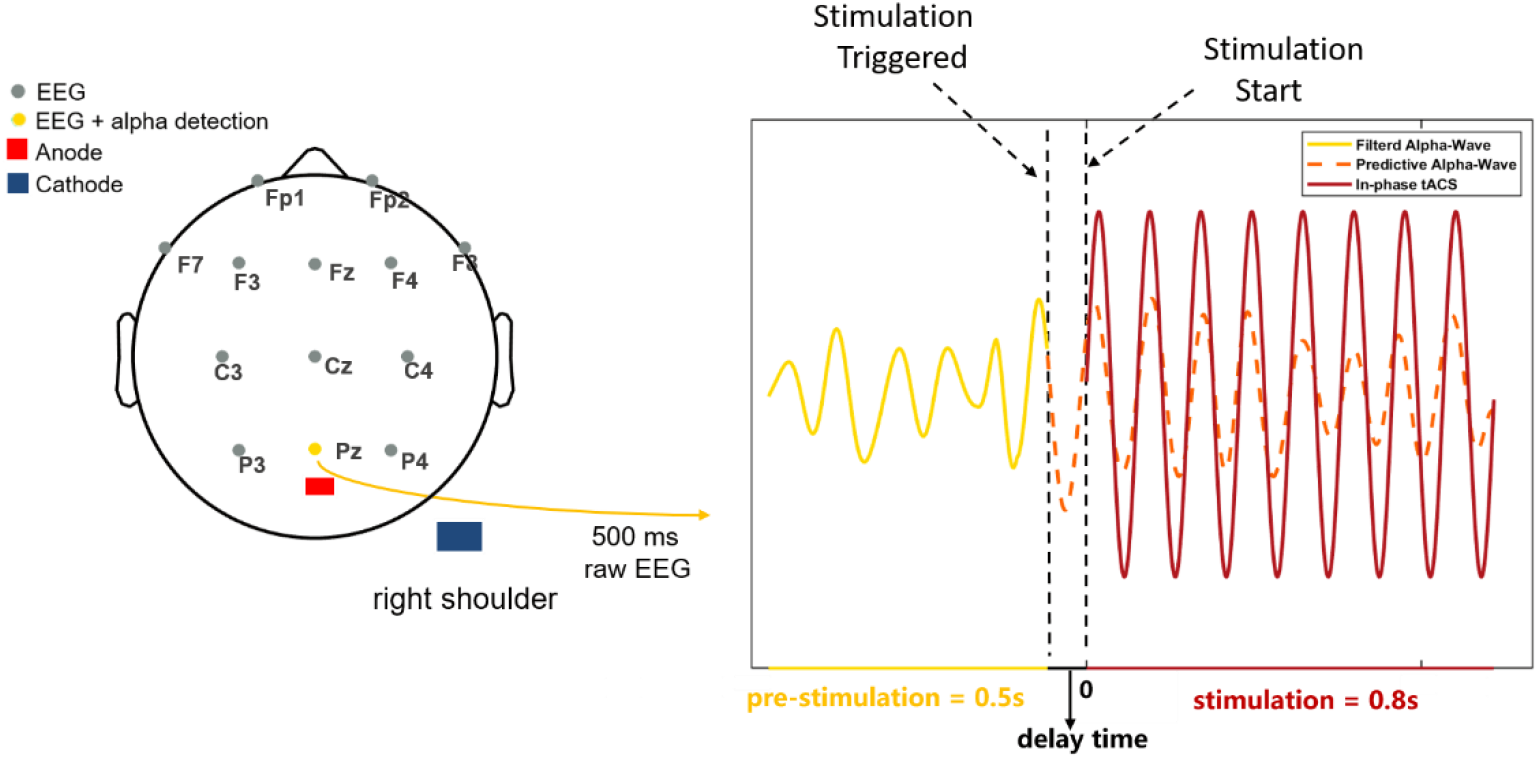
Illustration of the online phase-corrected closed-loop tACS system. The left panel shows a standard 10-20 electrode system used in the experiment; the right panel shows an example of alpha-wave detection and the application of in-phase tACS. The stimulation electrode (shown with the red rectangle in the left panel) was placed over the central parietal-occipital cortex (between the Pz electrode and Oz electrode), with the return electrode over the right shoulder (shown with the blue rectangle in the left panel). The raw EEG data at the retention interval of the Sternberg task, as measured from the Pz electrode, was stored in a moving time window of 500ms with a 10ms step. This signal was filtered within the frequency range of IAF±2Hz (shown with the dark-yellow solid line) in real time. When two consecutive peaks exceeding the threshold were detected (note that the threshold was determined using pre-test EEG data), stimulation was triggered; subsequently after a specific delay time, in-phase tACS (shown in dark red) was initiated and lasted for 0.8 seconds. The dark-orange dashed line shows the alpha wave after the triggering of stimulation (please note that it is not recorded due to tACS artifacts). The illustrative waveform in the figure does not represent the actual size.

**Fig. 3.**
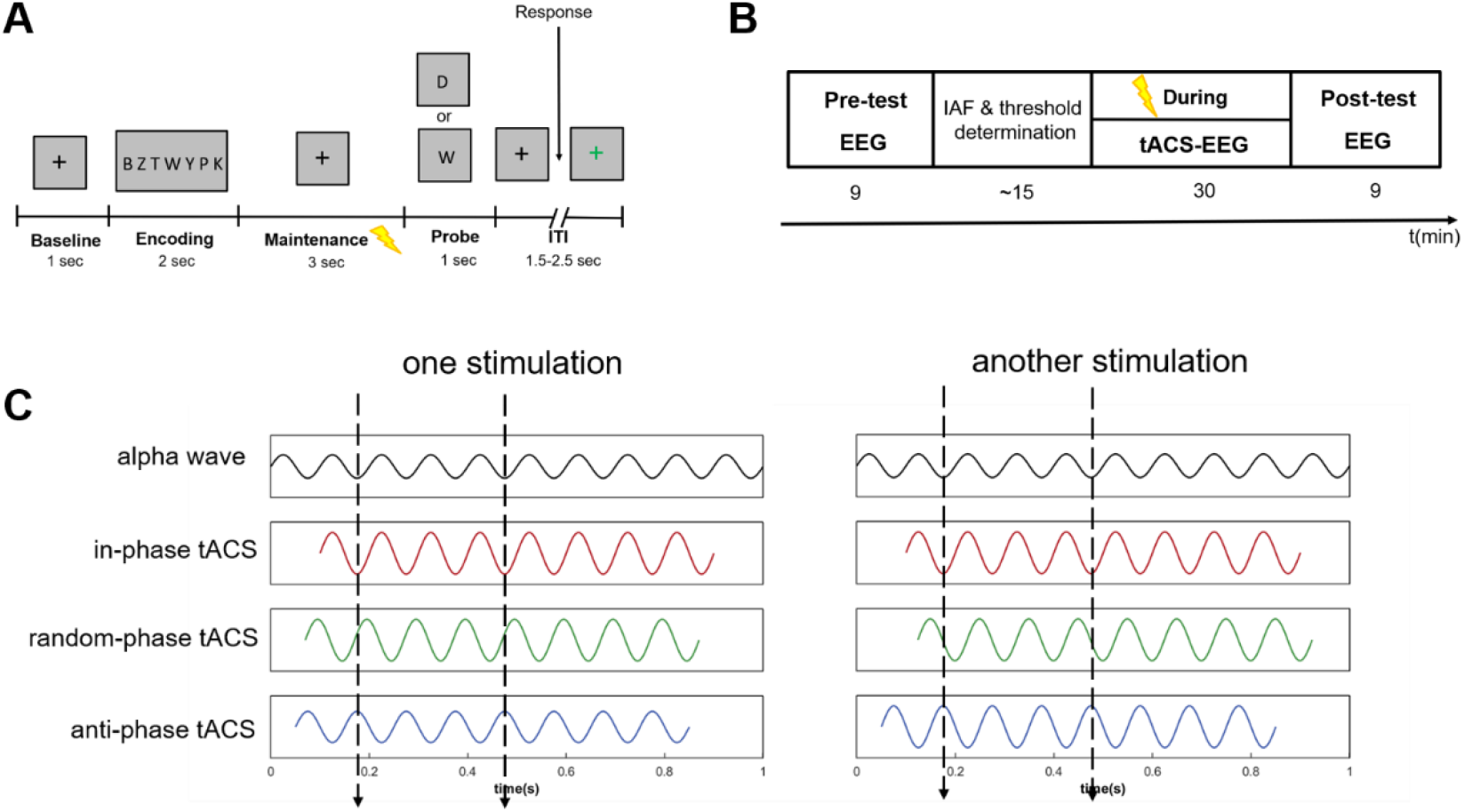
Sternberg task, experimental procedures, and stimulation conditions. (**A**) Schematic representation of the modified Sternberg paradigm used in this study. For each trial, participants were shown a list of 7 consonants and were asked to indicate by button press whether the probe was part of the memory list. (**B**) Experimental procedure for each session. First, participants completed a pre-test block of the Sternberg task with EEG recorded during the retention interval. Subsequently, the individual alpha frequency (IAF) of this session was determined using the pre-test EEG data. Then, 3 tACS-EEG blocks (During) of the Sternberg task were performed with tACS delivered specifically during the retention interval. After tACS, participants completed a final block of the Sternberg paradigm (post-test) while recording EEG signals. (**C**) Three tACS conditions: TACS was applied at the IAF with 0° relative phase to the endogenous alpha oscillations in the in-phase condition and with 180° relative phase to the endogenous alpha oscillations in the anti-phase condition; in the random-phase condition, no phase alignment between the delivered tACS and the detected endogenous alpha oscillations were used, with the phase differences changing from 0° to 360° across trials. For all conditions, the onset phase of the sine-wave tACS was invariably 0°. The illustrative waveforms in the figure do not represent the actual sizes.

## 2. Results

We used the online phase-corrected closed-loop tACS system to modulate alpha oscillations during the retention interval of WM (**Fig. 1** **& 2**; Methods; Supplementary material: 1.1. The details of online phase-corrected closed-loop tACS) while participants performed a Sternberg paradigm (**Fig. 3A**; Methods) [23]. TACS with 2-mA peak-to-peak amplitude at individual alpha frequency (IAF, see Supplementary material: 1.2. IAF and threshold determination) was applied to the central parietal-occipital brain areas in in-phase (with 0° relative phase to the endogenous alpha oscillations at Pz electrode), anti-phase (with 180° relative phase to the endogenous alpha oscillations at Pz electrode), and random-phase (the phase difference between the tACS stimulation and the endogenous alpha oscillations at Pz electrode was different across trials) conditions (**Fig. 3C**; Methods). Each participant performed a session of each tACS condition (in-phase tACS, anti-phase tACS, and random-phase tACS) separated by at least 3 days in a pseudo-randomized cross-over design. The experimental procedures for the three sessions are the same (**Fig. 3B**). To measure the behavioral performance of the Sternberg WM task, we calculated the rate correct score (RCS, see Methods for details on RCS), accuracy (defined as the percentage of correct responses), and mean reaction times (RT, only correct RTs were included in the analysis). The alpha power (8-13 Hz) at Pz electrode was calculated (see Methods for detailed information). The mean phase lag index (PLI) between the alpha activity (8-13Hz) of the 7 frontal electrodes (Fp1, Fp2, F7, F3, Fz, F4, F8) and Pz electrode was used to measure the frontoparietal alpha synchronization (see Methods for detailed information).

### 2.1. No systematic differences between in-phase tACS and anti-phase tACS at pre-test

In order to test the main hypothesis that tACS effects depend on the phase differences between tACS and endogenous brain oscillations, the different effects between in-phase tACS and anti-phase tACS were tested first. To ensure that there were no systematic differences between in-phase and anti-phase sessions before tACS, two-tailed paired *t*-tests were conducted for pre-test data. There were no significant differences in WM performance at pre-test for the two stimulation sessions (permuted paired *t*-test, accuracy: t(38) = 0.039, p = 0.956; RT: t(38) = −0.135, p = 0.894; RCS: t(38) = −0.002, p = 0.998). Further, there were no significant differences between the two sessions in pre-test alpha power or frontoparietal alpha synchronization values (permuted paired *t*-test, alpha power: t(38) = −0.316, p = 0.743; PLI: t(38) = −1.658, p = 0.104).

Participants performed the modified Sternberg task with high accuracy and fast reaction times, as is typical of young healthy participants: the mean accuracy across tACS conditions during pre-test is 0.857 (SD = 0.064), and the mean reaction time during pre-test is 878 ms (SD = 134 ms). These behavioral findings in our study are comparable to previous studies using this task [6].

### 2.2. Anti-phase tACS impaired working memory performance compared to in-phase tACS

Due to the suggested different mechanisms underlying online tACS effects [18, 25] (effects observed during stimulation: During) and offline tACS effects [26] (aftereffects beyond stimulation: post-test), we first analyzed the online tACS effects by comparing the online behavioral and EEG effects induced by in-phase and anti-phase tACS, then the offline tACS effects.

We first investigated whether anti-phase tACS down-regulated WM performance compared to in-phase tACS as hypothesized, measured by RCS, accuracy, and RT. At During, anti-phase tACS induced a significant RCS reduction compared to in-phase tACS (permuted paired *t*-test, t(38) = −2.209, p = 0.036, Cohen’s d = 0.354), which reflected fewer correct responses per second of activity in anti-phase tACS (**Fig. 4A**). Anti-phase tACS also induced a marginally significant reduction in accuracy when compared to in-phase tACS (permuted paired *t*-test, t(38) = −1.958, p = 0.057, Cohen’s d = 0.314) (**fig. S2A**). No significant difference in RT was observed between in-phase and anti-phase tACS at During (**fig. S2B**) (See more details in Supplementary material: 2. The online comparison of accuracy and RT between in-phase tACS and anti-phase tACS).

**Fig. 4.**
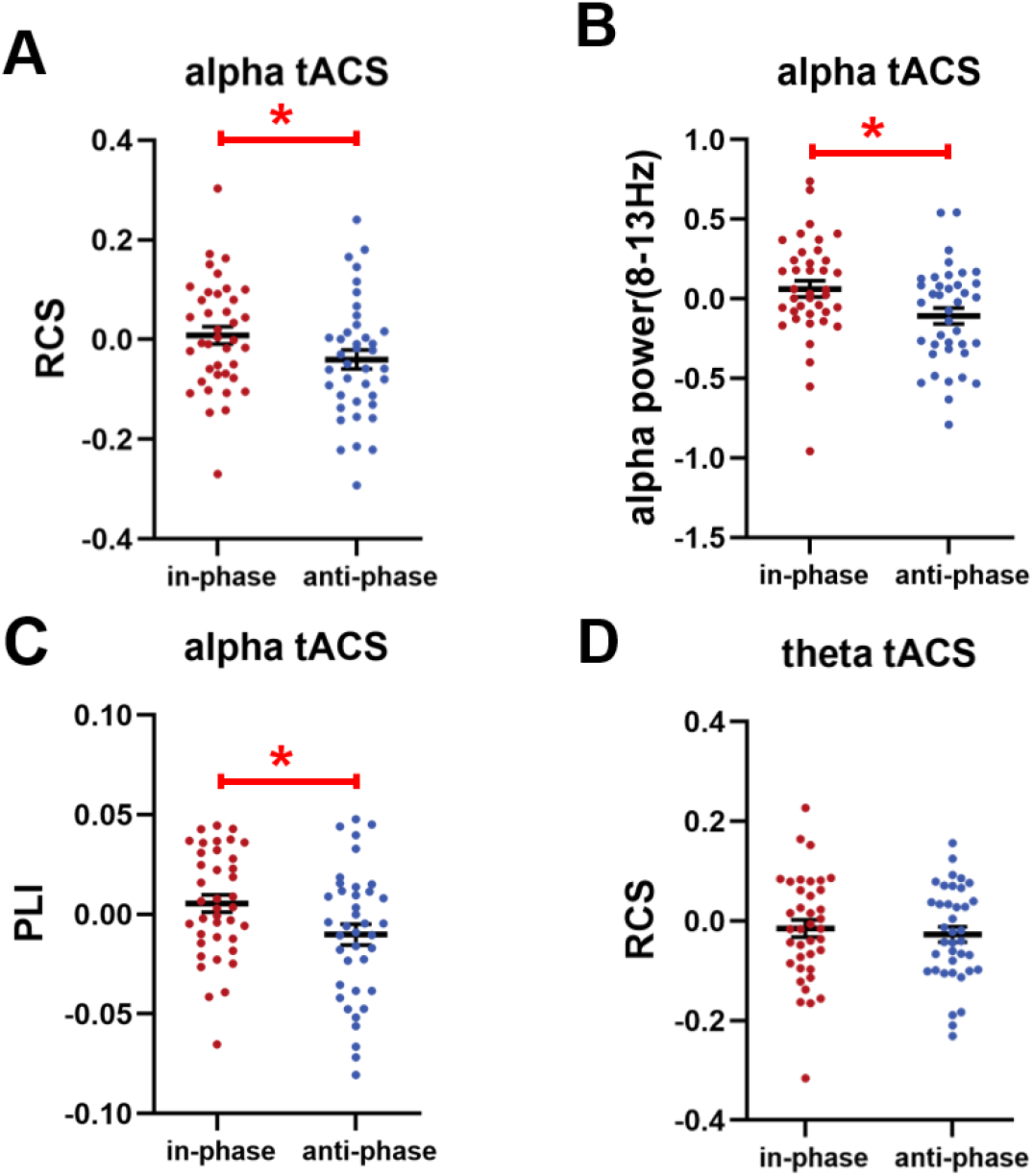
Compared to in-phase tACS, anti-phase tACS at the alpha frequency band decreased alpha activity and WM performance. (**A**)Effects of anti-phase tACS delivered at the IAF on the rate correct score (RCS, the number of correct responses per second). At During, anti-phase tACS significantly decreased the RCS as compared to in-phase tACS (permuted paired *t*-test: t(38) = −2.209, p = 0.036). (**B**) Anti-phase tACS at the IAF significantly suppressed parietal alpha power (8-13 Hz) as compared to in-phase tACS at During (permuted paired *t*-test: t(38) =-2.257, p = 0.027). (**C**) As compared to in-phase tACS, frontoparietal alpha synchronization in anti-phase tACS (indexed by PLI) decreased significantly at During (permuted paired *t*-test: t(38) =- 2.067, p = 0.044). (**D**) For tACS at the theta frequency band, no suppression effects on WM performance were detected between the anti-phase tACS and in-phase tACS. Note that all instances of the RCS, alpha power, and frontoparietal alpha synchronization values are given relative to the pre-test data (i.e., values after subtracting the corresponding pre-test values). Within-group comparisons used two-tailed permuted paired t-tests. Error bars represent the SEM; *significant at p <0.05 (two-tailed permuted paired *t*-tests), ** significant at p <0.01 (two-tailed permuted paired *t*-tests).

### 2.3. Anti-phase tACS suppressed alpha power compared to in-phase tACS

We next assessed whether in-phase tACS and anti-phase tACS modulated the targeted parietal alpha oscillations during stimulation. Alpha power (8-13Hz) at Pz electrode was analyzed because its signals served for triggering tACS and it was near to the stimulation electrode. As hypothesized, anti-phase tACS significantly suppressed the alpha power at Pz electrode compared to in-phase tACS at During (permuted paired *t*-test, t(38) =-2.257, p = 0.027, Cohen’s d = 0.361) (**Fig. 4B**). This finding suggests the modulation effects of the closed-loop tACS system on the targeted brain activity and the influence of phase differences on tACS electrophysiological effects.

To support the frequency-specific effects of the online phase-corrected closed-loop tACS system, we next tested the influences of alpha tACS on the full physiological frequency band of the EEG (1-45 Hz). No significant differences between in-phase tACS and anti-phase tACS were found in any of the frequency bands except for the alpha band at During (For more details, see Supplementary material: 3.1 The comparison of power in full frequency band between in-phase alpha-tACS and anti-phase alpha-tACS). This result indicates the frequency-specific modulation of phase-corrected closed-loop alpha tACS on endogenous brain oscillations as previous tACS studies have shown [12, 26] (**fig. S3**). Moreover, we also calculated the modulation effects of alpha-tACS on individual alpha power (IAF ± 2Hz) at Pz electrode, and found that anti-phase tACS also showed suppression effects on individual alpha power (**fig. S4**), further supporting the modulation effects of alpha-tACS on parietal alpha oscillations. Therefore, it is reasonable to attribute the behavioral effects of tACS to the modulation on alpha oscillations at the retention interval, supporting the causal link between alpha oscillations and WM retention.

### 2.4. Anti-phase tACS disturbed frontoparietal alpha synchronization compared to in-phase tACS

Previous studies have reported the potential impact of tACS on functional connectivity [27, 28], and found increases in the strength of alpha synchronization with increasing memory load among the frontoparietal regions known to underlie executive and attentional functions during WM maintenance [29, 30]. Therefore, online tACS effects in frontoparietal alpha synchronization were also assessed.

Anti-phase tACS induced a significant decrease in frontoparietal alpha synchronization compared with in-phase tACS at During (permuted paired *t*-test, t(38) =-2.067, p = 0.044, Cohen’s d = 0.331) (**Fig. 4C**). To further illustrate the reliability of our results, we also used weighted phase lag index (WPLI) to measure frontoparietal alpha synchronization, which may be more sensitive to unrelated noise resources [31]. As expected, compared to in-phase tACS, anti-phase tACS disturbed frontoparietal alpha synchronization, measured by WPLI, at During (**fig. S5)**. These results suggest that the online phase-corrected closed-loop tACS not only affect the brain activity of the targeted region, but also modulate the connectivity of distributed brain regions.

Besides, there were no significant differences between tACS conditions (in-phase and anti-phase) for tACS-induced side effects (for more details of the analysis, see the Supplementary material: 1.4. Analysis of tACS questionnaire), so it rules out that the above reported different behavioral and electrophysiological effects between in-phase tACS and anti-phase tACS were caused by tACS-related sensations.

### 2.5. Compared to pre-test, anti-phase tACS significantly down-regulated WM performance and alpha power at the stimulation period

Above results showed that compared to in-phase tACS, anti-phase tACS induced a significant online down-regulation in WM performance, alpha power, as well as frontoparietal alpha synchronization at During. To further investigate whether the down-regulation was due to in-phase tACS-induced improvement or anti-phase tACS-induced suppression, we analyzed the changes from pre-test to During within in-phase tACS and anti-phase tACS. RCS, accuracy, and alpha power at Pz electrode in anti-phase tACS reduced significantly at During (permuted paired *t*-test, RCS: t(38) = −2.117, p = 0.047, Cohen’s d = 0.339; accuracy: t(38) = −2.434, p = 0.019, Cohen’s d = 0.390; alpha power: t(38) = −2.171, p = 0.036, Cohen’s d = 0.348). Anti-phase tACS also induced marginally significant disturbance in frontoparietal alpha synchronization at During (permuted paired *t*-test, t(38) = −1.926, p = 0.061, Cohen’s d = 0.308). For in-phase tACS, no significant improvement was observed from pre-test to During for all of the metrics (permuted paired *t*-test, RCS: t(38) = 0.467, p = 0.642; accuracy: t(38) = −0.152, p = 0.887; RT: t(38) = −0.118, p = 0.909; alpha power: t(38) = 1.223, p = 0.225; PLI: t(38) = 1.266, p = 0.207); the reasons for the non-significant enhancement effects of in-phase tACS may be due to the difficulty to further enhance endogenous strong oscillations by intermittent tACS (See Supplementary material: 8.2. A discussion on the non-significance of in-phase tACS-induced enhancement effects). These results indicated that the different effects between in-phase tACS and anti-phase tACS at During were mainly due to the suppression effects of anti-phase tACS.

### 2.6. The correlations between tACS-induced changes in alpha activity and RCS were significantly positive for in-phase condition, which was different from anti-phase condition

As alpha-tACS not only altered RCS, but also modulated alpha power and frontoparietal alpha synchronization at During, we further investigated whether the modulation of alpha activity related to WM performance at During. We performed permuted Pearson’ s correlations between the stimulation-induced changes (from pre-test to During) in RCS and EEG metrics. The correlation between in-phase tACS-induced changes in alpha power and RCS was significant at During (permuted Pearson’s correlation, r = 0.348, p = 0.030). This finding indicated that across participants, increased alpha power related to larger RCS, further supporting the causal role for alpha oscillations in WM retention. No correlation was found between anti-phase tACS-induced changes in alpha power and RCS (permuted Pearson’s correlation, r = −0.244, p = 0.138), possibly because anti-phase tACS disrupted the inherent relationships between parietal alpha oscillations and WM performance in the absence of exogenous disturbance. Notably, there was a significant difference between the correlation coefficients in in-phase tACS and anti-phase tACS at During (Z = 2.599, p = 0.009) (**Fig. 5A**).

**Fig. 5.**
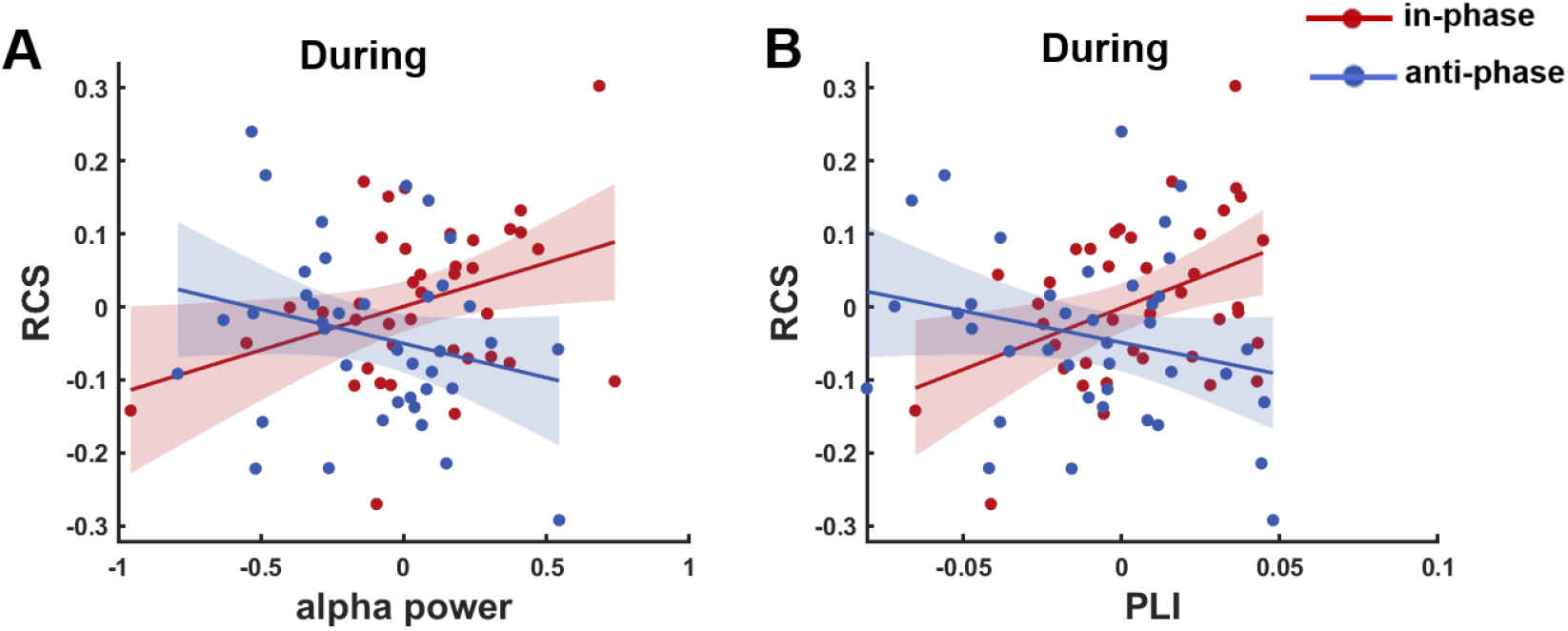
Correlations between tACS-induced changes in RCS and EEG metrics differed between in-phase tACS and anti-phase tACS. (**A**) Permuted Pearson’s correlation (two-tailed) analysis was used to assess the potential relationship between RCS and parietal alpha power at During: the in-phase tACS-induced changes in RCS was positively correlated with the induced changes in alpha power (permuted Pearson’ correlation: r = 0.348, p = 0.030); no correlation was found for anti-phase tACS (permuted Pearson’ correlation: r = −0.244, p = 0.138). There was a significant difference between the correlation coefficients in in-phase tACS and anti-phase tACS (Z = 2.599, p = 0.009). (**B**) Similar assessment of the relationship between tACS-induced changes in RCS and induced changes in frontoparietal alpha synchronization at During. A positive correlation between the tACS-induced changes in frontoparietal alpha synchronization and RCS was found for in-phase tACS (permuted Pearson’ correlation: r = 0.415, p = 0.007) but not for anti-phase tACS (permuted Pearson’ correlation: r = −0.242, p = 0.140). These two correlation coefficients differed significantly between in-phase tACS and anti-phase tACS (Z = 2.921, p = 0.003). The RCS, alpha power, and frontoparietal alpha synchronization values are given relative to the pre-test values (i.e., after subtracting corresponding pre-test values). The obtained permuted Pearson’s correlation coefficients were converted into z-values using Fisher z-transformation to support comparison of two correlations.

Similar to the correlations between the changes in alpha power and RCS, we also found a positive correlation between tACS-induced changes in frontoparietal alpha synchronization and RCS at During for in-phase tACS (permuted Pearson’s correlation, r = 0.415, p = 0.007), and no correlation for anti-phase tACS (permuted Pearson’s correlation, r = −0.242, p = 0.140). A significant difference between the correlation coefficients in in-phase tACS and anti-phase tACS was also observed at During (Z = 2.921, p = 0.003) (**Fig. 5B**). The observed inverse correlation trends for in-phase tACS and anti-phase tACS further illustrate the different effects between in-phase tACS and anti-phase tACS on WM.

### 2.7. The effects of random-phase tACS and the offline tACS effects

To explore the effects of the control condition, here we compared the effects of random-phase tACS with both in-phase and anti-phase tACS. At During, random-phase tACS-induced changes in WM performance (**Fig. 6A**), the alpha power at Pz electrode (**Fig. 6B**) and frontoparietal alpha synchronization (**Fig. 6C**) were all between in-phase tACS and anti-phase tACS. The correlations between random-phase tACS-induced changes in alpha activity and RCS were not significant (**Fig. 6D** **&** **E**), also between in-phase tACS and anti-phase tACS (See the detailed results for random-phase tACS in Supplementary material: 5. Complementary analysis of the behavior and EEG data in random-phase tACS). These results indicate the effects of random-phase tACS were between in-phase tACS and anti-phase tACS, further supporting the influence of the phase differences on tACS effects.

**Fig. 6.**
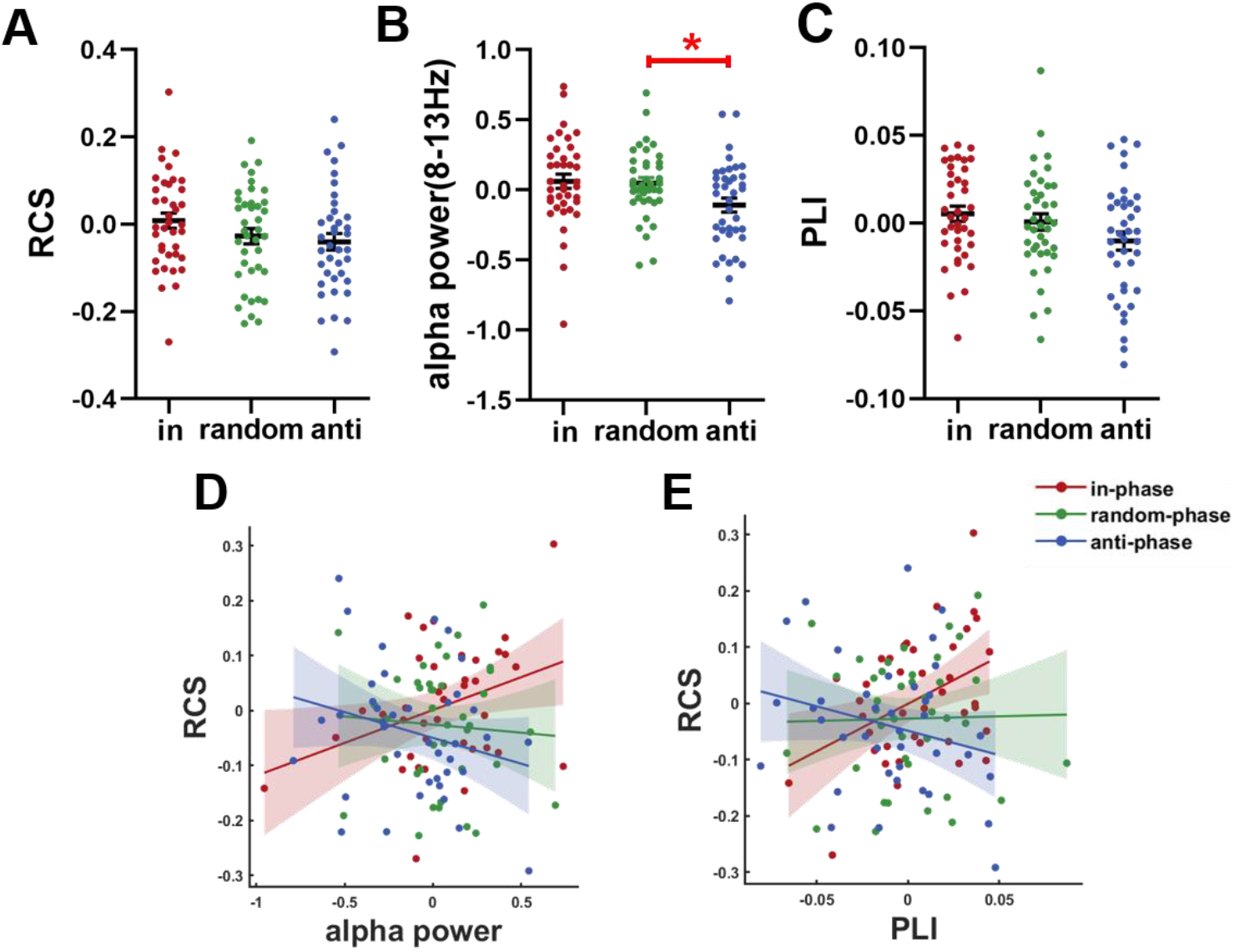
At During, the effects of random-phase tACS were between in-phase tACS and anti-phase tACS. Random-phase tACS-induced changes in (**A**) RCS, (**B**) alpha power at Pz electrode, and (**C**) frontoparietal alpha synchronization were all between in-phase tACS and anti-phase tACS. The random-phase tACS-induced changes in RCS were not significantly correlated with the induced changes in (**D**) alpha power and (**E**) frontoparietal alpha synchronization. The RCS, alpha power, and frontoparietal alpha synchronization values are given relative to the pre-test values (i.e., after subtracting corresponding pre-test values). Error bars represent the SEM; *significant at p <0.05 (two-tailed permuted paired *t*-tests).

Next, we explored the offline tACS effects (tACS effects beyond the stimulation period: post-test) on WM performance and brain activity. We observed no significant difference between in-phase tACS and anti-phase tACS in WM performance, alpha power at Pz electrode, and frontoparietal alpha synchronization (**fig. S9**) (For more details about offline effects, see Supplementary material: 6. The offline comparison between in-phase tACS effects and anti-phase tACS effects).

### 2.8. Frequency-specific effects of tACS on WM performance

To provide additional support to the conclusion that the observed changes in WM performance can be attributed to changes in parietal alpha oscillations during the WM retention interval, rather than to some general effects from tACS (independent of frequency) [24], we conducted a control experiment in which both in-phase tACS and anti-phase tACS were delivered in the theta frequency band (3-8Hz) to 38 participants (For more details about theta-tACS experiment, see Supplementary material: 7. In-phase and anti-phase tACS at the theta frequency). No suppression effects from the anti-phase theta-tACS were observed on WM performance (RCS) at any time point (During, post-test) as compared to the in-phase theta-tACS (permuted paired *t*-test, During: t(37) = −0.512, p = 0.617; post-test: t(37) = 0.334, p = 0.736) (**Fig. 4D**). This lack of any detected impact from theta-tACS supports that modulation of alpha oscillations—rather than some general impacts from electrical stimulation—can explain the observed effects of alpha-tACS on WM performance.

## 3. Discussion

In this study, we used an online phase-corrected closed-loop tACS system to modulate alpha oscillations specifically during the WM retention interval on a trial-by-trial level, with a predetermined phase difference between the tACS and the concurrent endogenous oscillatory activity. Compared to in-phase tACS, anti-phase tACS decreased working memory performance, and these decreases were paired with corresponding decreases in alpha power and in frontoparietal alpha synchronization at the stimulation period. Notably, the detected changes in alpha power and frontoparietal alpha synchronization induced by in-phase tACS were both positively correlated to behavioral changes. Our study therefore provides direct causal evidence of a specific functional impact of alpha oscillatory activity in human WM retention. Our work also illustrates that phase differences between tACS and the target brain oscillations represent a decisive factor for determining the effects of tACS, both on neural modulation and on behavioral performances.

### 3.1. Causal role for alpha oscillations in the retention sub-process of WM

Our behavioral and electrophysiological findings strongly support a causal link between parietal alpha oscillations and WM retention. Alpha oscillations during the memory retention interval have been suggested to be instrumental in transiently protecting the encoded memory information by filtering task-irrelevant input and preventing further sensory processing that could interfere with the stored information [5, 7, 32, 33]. However, no direct causal evidence supporting this functional role of alpha oscillatory activity has been established in human neuroscience to date. A few previous studies using non-invasive brain stimulation (NIBS) attempted to investigate alpha oscillations and WM retention in humans [34–36], but multiple aspects of these studies prevent causal inferences. Due to the incapacity of previous tACS stimulation to be time-locked to the short-time sub-process of WM, one alpha-tACS study applied continuous tACS throughout the three sub-processes of WM rather than specifically during retention interval, so it was not sufficient to establish a functional role of alpha oscillations in WM retention *per se* [34]. rTMS studies tried to modulate neural oscillations during the retention interval, but did not measure rTMS-induced changes in endogenous alpha activity, so the attribution of the observed behavioral effects on WM performance to the modulation of alpha oscillations remains speculative [35, 36]. Addressing these aspects, we used our online phase-corrected closed-loop tACS system to modulate parietal alpha oscillations specifically during WM retention, while also recording EEG during intervals without tACS artifacts to investigate the influence of external modulation on alpha oscillations. We found that anti-phase tACS in our study induced both WM performance impairment and parietal alpha inhibition at the stimulation period. We also observed a positive association between in-phase tACS-induced changes in parietal alpha power and the changes in working memory performance, further supporting the causal link between alpha activity and WM retention.

### 3.2. Frontoparietal alpha synchronization contributes to WM performance

Our findings also suggest that frontoparietal alpha synchronization contributes to WM. Anti-phase tACS induced a significant decrease in frontoparietal alpha synchronization at the stimulation period. This decrease may be attributed to disturbance of the parietal alpha phases by anti-phase tACS, which likely impairs its coupling with frontal brain areas. A few studies have found that synchrony is strengthened with increasing memory load in frontoparietal regions previously shown to mediate attentional functions during memory retention [29, 30]. In line with these studies, we observed a positive association between in-phase tACS-induced changes in frontoparietal alpha synchronization and altered RCS values, which demonstrates that increased frontoparietal alpha synchronization during WM retention also contributes to the observed improvement in WM performance.

### 3.3. The phase differences between tACS and endogenous brain oscillations affect tACS effects

In agreement with previous computational modelling predictions [18] and animal study [19] suggesting that tACS with different phases relative to endogenous brain oscillations would induce different modulation effects on electrophysiological activities, we experimentally demonstrated that anti-phase tACS significantly inhibited parietal alpha activity, frontoparietal alpha synchronization, and working memory performance compared to in-phase tACS at the stimulation period. Therefore, we provide the direct experimental evidence that the phase of endogenous brain oscillations relative to tACS is impactful for determining the direction and magnitude of tACS. This evidence also calls for the consideration of brain states during tACS (e.g., the phase differences between the tACS waveform and the endogenous alpha oscillations) to overcome the unwelcome inconsistent effects of conventional open-loop tACS [17]. The suppression effects of anti-phase tACS compared to in-phase tACS also suggest that tACS can act by the modulation of cortical neurons, rather than through peripheral nerve stimulation in the scalp [37]. Effects from in-phase tACS vs. anti-phase tACS should not differ if they only result from transcutaneous stimulation [37] based on electrodes positioned at the head and the right shoulder.

### 3.4. The advantages and application potentials of the online phase-corrected closed-loop tACS

The capacity to induce or suppress target oscillatory activities within a relatively short period of time (within several seconds) is a unique and highly valuable feature of our online phase-corrected closed-loop tACS system as compared to other currently available tACS techniques. Conventional continuous tACS was usually demonstrated to enhance brain oscillations at the stimulation frequency via entrainment [13]. Although several studies reported the suppression effects of conventional long-term continuous tACS on brain oscillations at the non-stimulation frequency during specific brain states [38, 39] (e.g., the suppression effects of alpha tACS on gamma oscillations in a visual detection task [39]), this kind of inhibition effects cannot be generalized because the correspondence between the tACS frequency and the frequencies of the suppressed brain oscillations is not clear. Moreover, recently published brain-signal based short-time closed-loop tACS set-ups have to date only reported unidirectional modulation of brain oscillations at the stimulation frequency [40–43]. This inflexibility regarding directionality of tACS effects within a short period of time is problematic for many functions and applications. Consider for example that previous studies of the Sternberg paradigm have indicated that alpha activity tends to increase during the retention sub-process but to decrease during the encoding and retrieval sub-processes [21, 22]. If conventional long-term (several to dozens of minutes) continuous tACS was performed throughout the entire task process, the effects in different sub-processes might cancel each other out. We selectively regulated alpha oscillations during the retention sub-process; future studies could try to up-regulate alpha activity during the retention sub-process while also—and in the same trial—down-regulating oscillatory alpha activity or even manipulating related oscillations at other frequencies during the encoding and retrieval sub-processes. This method, or many variations thereof, may prove to be an even more effective way of modulating cognitive processes, e.g., working memory.

It is further plausible to speculate that our phase-corrected closed-loop tACS system may broaden the clinical therapy applications of tACS, with its expanding capacity to suppress brain oscillations within a relatively short period. Our results show that anti-phase tACS can significantly inhibit oscillatory activity, illustrating the possibility of using tACS to down-regulate abnormally high brain rhythms, especially those excessive brain oscillations that only occur within a short time during the attacks of diseases. For example, our anti-phase tACS may be suitable to suppress the cue-induced excessive low-frequency oscillations (e.g., increased delta or theta oscillations) of cocaine addicts within a short period [44], which cannot be achieved by conventional open-loop tACS without considering the phase differences.

### 3.5. Conclusion

In summary, our study provides empirical evidence for a causal link between parietal alpha oscillations and the retention sub-process of working memory. To establish this link, we here pioneered an online phase-corrected closed-loop tACS system that offers the possibility of directly up- and down-regulating oscillatory brain activities at a predetermined stimulation frequency, and we show that this system does induce effects on both brain rhythms and behavior.

## 4. Materials and Methods

### 4.1. Participants

We calculated the needed sample size using G*Power 3.1 software [45]. The main purposes of the present study were to investigate whether anti-phase alpha-tACS and in-phase alpha-tACS would induce different behavioral and electrophysiological effects, and whether changes in WM performance were associated with changes in alpha activity. Assuming a medium effect size (Cohen’s d = 0.5) for two-tailed paired t-tests with a statistical power of 80% and an alpha probability of 0.05, a sample size of 34 participants were needed. Assuming a correlation coefficient of medium effect size (r = 0.4) and a statistical power of 80%, a sample size of 44 participants were needed.

A total of 48 healthy participants between the ages of 18 and 40 completed the alpha-tACS experiment. All participants were right-handed, reported no metal implants in the brain, no implanted electronic devices, no history of neurological problems or head injury, no current use of psychoactive medication, no history of craniotomy, no skin sensitivity, were nonpregnant, had normal or corrected-to-normal visual acuity, and not being enrolled in any other NIBS research within 3 months of their study participation. Participants who failed to follow directions or did not understand instructions were removed from the study. All participants were recruited in Hefei, China through online advertisements or posters. Eight participants were excluded from the analyses due to poor EEG signals or malfunctions of the tACS stimulator to trigger intended electrical stimulation. One participant whose alpha activity was too weak for detection and who therefore had to repeat the pre-test Sternberg paradigm 3 times per session was also excluded. The remaining 39 participants (19 females, mean age ± SD: 21.1 ± 2.2 years, mean education ± SD: 14.7 ± 1.8 years) were tested and their data were submitted to behavioral and EEG analyses, which meets the sample size requirement.

The study was approved by the Human Ethics Committee of the University of Science and Technology of China (IRB No.: 2020KY161) and performed in accordance with the latest Declaration of Helsinki. All participants provided written informed consent prior to the study and were paid 300 RMB after completing three experimental visits.

### 4.2. Experimental procedures

Each participant underwent 3 experimental sessions separated by at least 3 days at approximately the same time of the day. Using a single-blinded within-subject design, participants received in-phase tACS, anti-phase tACS, or random-phase tACS in each session (**Fig. 3C**), with the order of tACS conditions counterbalanced across participants.

The experimental procedures for the three sessions are the same (**Fig. 3B**). At the start of each session, preparing the closed-loop tACS-EEG setup required about 30 minutes per participant. Following this preparation period, participants first completed a practice block of the Sternberg paradigm (**Fig. 3A**) consisting of 32 trials. The Sternberg task is chosen due to its suitability to separate the three different sub-processes of working memory: encoding, retention, and retrieval, so that tACS can be applied specifically during the retention interval on a trial-by-trial basis. Following this practice, participants performed 5 blocks of the Sternberg paradigm. Within each block, there were 30 positive trials (probe part of the memory list) and 30 negative trials (probe not part of the memory list).

In the first block (pre-test), spontaneous EEG was recorded for 2.5s in each trial once the retention interval of Sternberg task began. Subsequently, the EEG data at Pz electrode of the pre-test block was analyzed to determine the IAF to be used as the subsequent tACS frequency for this session, and the threshold for subsequent triggering of tACS (see more details in Supplementary material: 1.2. IAF and threshold determination). The threshold was set to really target the interested endogenous oscillations (i.e., alpha oscillations) by delivering tACS only when strong endogenous interested oscillations exist.

Then, participants performed 3 tACS-EEG blocks (During) while performing the Sternberg task. EEG data at Pz electrode was monitored once the retention interval began. The closed-loop stimulation was triggered and lasted for 0.8 s when the EEG signal met requirements (for detailed description of the closed-loop tACS, see Online phase-corrected closed-loop tACS section below and Supplementary material: 1.1. The details of online phase-corrected closed-loop tACS). The 0.8s duration of each stimulation was a balance of total effective stimulation time and the consistency of the phase differences between brain signals and tACS which has been shown to decrease as the length of the tACS time window increases [46]. Once stimulation started, the EEG recording stopped to avoid tACS artifacts until the onset of the next trial.

After a rest period of about 2 min, participants completed a second WM task for about 30 min. The results of this part will be reported in other places. After the second WM task, participants filled out a tACS questionnaire to assess tACS-induced discomfort (see more details in tACS sensations section). Finally, participants completed a final block of the Sternberg paradigm (post-test) while recording EEG.

### 4.3. Sternberg task

Participants performed a previously published modified version of the Sternberg WM task [6, 8] (**Fig. 3A**). Participants had to remember a horizontally arranged list of seven consonants that was presented simultaneously at the center of the computer monitor for 2 s. After a 3 s retention interval (blank screen with fixation cross on the middle position of the monitor), the probe stimuli were presented and displayed for the duration of the recognition interval (1 s). All responses were made with the right hand, by pressing the Left arrow with the index finger and the Right arrow with the middle finger. Participants had to press a button if the probe matched one of the consonants in the memory set, and press the other button if the probe did not match. Instructions stressed both speed and accuracy. The inter-trial interval (ITI) was a uniformly distributed random number between 1.5s and 2.5s. A fixation cross was presented during the ITI. During the trial, the letters and the fixation cross were presented in black on a gray background. After the button press, a green fixation cross was shown, indicating that the participant responded successfully and was allowed to make eye blinks. One second before the start of the next trial the fixation cross turned black again. Participants were instructed not to blink from this moment until they had pressed a button.

### 4.4. EEG data collection

EEG data was recorded using a UEA-16BZ amplifier (SYMTOP, Beijing, China). Thirteen Ag/AgCl electrodes were placed on the scalp at specific locations according to the international 10-20 system (Fp1, Fp2, F7, F3, Fz, F4, F8, C3, Cz, C4, P3, Pz, P4). In addition, the electrical activities were recorded over left and right mastoids. As previous studies reporting the relationship between alpha oscillations and WM retention usually used mastoids as reference [6, 22], the average activities of bilateral mastoids were used as reference in this study. Impedance between the reference and any recording electrode was kept under 5 kΩ. All signals were sampled at 1000 Hz during data collection. A low-pass filter with a cut-off frequency of 45Hz and a 50Hz notch filter were applied on-line.

For the pre-test and post-test blocks without tACS, EEG recording started from the onset of the memory retention sub-process and lasted for 2.5 s in each trial. At During, EEG signal in each trial was also recorded from the onset of the retention sub-process, but EEG recording stopped once tACS was triggered to avoid amplifier saturation and potential tACS artifacts considering the very close distance between tACS stimulation electrode and Pz electrode. In the trials where tACS was not triggered, EEG recording lasted for 2.5 s since the beginning of the retention sub-process.

### 4.5. Online phase-corrected closed-loop transcranial alternating current stimulation (tACS)

In order to be able to stimulate specifically at the retention interval in each trial with a predetermined phase difference between the applied stimulation and the endogenous oscillatory activities at the target brain region, we here introduced an online phase-corrected closed-loop tACS setup pioneered by our research group. The online phase-corrected closed-loop tACS system is able to measure brain oscillations by an EEG instrument, analyze the raw EEG data online using the computer to extract the phases of an underlying brain rhythm, control the timing of tACS based on the phases of the underlying oscillations, and closing the loop, affect the brain oscillations (**Fig. 1**). As shown in **Fig. 2**, The system enabled us to administer sinusoidal current stimulation with a certain phase difference relative to the target brain oscillations (for more details about how to realize the certain phase differences between tACS and brain oscillations, see the Supplementary material: 1.1. The details of online phase-corrected closed-loop tACS).

Then we used the online phase-corrected closed-loop tACS to provide 0.8 s sinusoidal alternating current stimulation of 2 mA (peak to peak) amplitude with 0° (in-phase tACS) or 180 ° (anti-phase tACS) phase difference relative to alpha oscillations (IAF ± 2Hz) recorded from Pz electrode. The activities of Pz electrode were chosen to be monitored because Pz electrode was reported to exhibit strongest alpha power during the retention interval of Sternberg paradigm[6]. The stimulation electrode (4×6 cm, rubber electrode covered with conductive paste) was placed over central parietal-occipital cortex (between Pz and Oz, according to the international 10-20 system), with the upper edge about 2 cm away and the center about 4 cm away from Pz electrode, to prevent the inevitable electrical noises produced by the standby-mode stimulator from contaminating EEG data at Pz electrode. Although the stimulation electrode was not placed precisely over Pz electrode, the position of it in our study is suitable due to the high coherencies of EEG signal at about 5 cm electrode separations [47] and the relatively large sources accounting for alpha activity in WM, extending from parietal region to occipital region [8] (Supplementary material: 8.1. A discussion on the position of the stimulation electrode). The return electrode (6×9 cm, rubber electrode covered with conductive paste) was over the right shoulder. Ten20 conductive gel (Nuprep, Weaver and Company, Aurora, CO, USA) was applied on the electrodes to reduce the impedances of each electrode to below 10 kΩ and hold the electrodes in place. Then, a custom 13-channel EEG cap (SYMTOP, Beijing, China) was worn by the participants on top of the stimulation electrodes (See EEG data collection).

The phase-corrected closed-loop tACS system ran for about 30 min during the 3 blocks of During; no stimulation was delivered during the first (pre-test) or final block (post-test). In order to explore the role of alpha oscillations in the retention sub-process, the closed-loop tACS system only monitored alpha oscillations and applied tACS during the retention interval of each trial. In order to avoid the duration of electrical stimulation beyond the retention interval of WM, monitoring for triggering the electrical stimulation was limited to 1.8 s since the onset of the memory retention interval. If the electrical stimulation was not triggered within 1.8 s, the system stopped monitoring alpha oscillations and continued recording the EEG data to 2.5s. During the running of the phase-corrected closed-loop tACS system, the phases of alpha oscillations at Pz electrode in a moving 500 ms time window with 10 ms step were extracted online by the computation module to provide the timing of tACS and the timing was communicated to the tACS stimulator (see the Supplementary material: 1.1. The details of online phase-corrected closed-loop tACS). Then, the stimulator was set to provide a sinusoidal current stimulation at IAF lasting for 0.8 s (IAF was calculated using the EEG signal of the first block (pre-test), see more details in Supplementary material: 1.2. IAF and threshold determination). The onset phase of the sine-wave stimulation was invariably 0 in all conditions. To ensure the real-time phase correction, there were no ramp up and ramp down periods in the 0.8 s stimulation. Any transmission delays in the process were accounted for (See more details in Supplementary material: 1.1. The details of online phase-corrected closed-loop tACS).

To test the accuracy of the correction for the phase differences, we analyzed the phase alignment between unstimulated EEG signals recorded at pre-test and artificial tACS waveform predicted offline, and the phase alignment is as expected (see Supplementary material: 1.3. Analysis of the phase alignment between EEG signal and tACS waveform for detailed information).

### 4.6. TACS sensations

Before the During period of each session, all participants were exposed to tACS stimulation for several times (no more than 5 times, 0.8 s each time) to make sure that the tACS with 2 mA amplitude was acceptable for participants. One participant who reported relatively severe painful sensations during the exposure was terminated from participation and released. After tACS finished, participants completed a questionnaire to assess possible side-effects of the tACS stimulation (see the Supplementary material: 1.4. Analysis of tACS questionnaire).

Analysis of the tACS questionnaires indicated that there were no significant differences between tACS conditions (in-phase, random-phase, and anti-phase) for tACS-induced side effects (for more details of the analysis, see the Supplementary material: 1.4. Analysis of tACS questionnaire). All participants confirmed that the stimulation was acceptable during tACS, and did not induce severe discomfort after the experiment.

### 4.7. Blinding

Participants were blinded to the stimulation conditions. Participants invariably received verum stimulation of the same amplitude (2 mA, peak to peak) in all of the three tACS conditions (i.e., without sham stimulation). Moreover, self-reported questionnaires for sensations elicited by tACS didn’t differ between conditions (for more details, see the Supplementary material: 1.4. Analysis of tACS questionnaire), further indicating that participants cannot distinguish between different stimulation conditions.

### 4.8. Offline EEG analysis

Analysis of the EEG data was performed using custom-built scripts implemented in MATLAB 2016a (MathWorks, Natick/USA) using the EEGLAB Toolbox (version 13.5.4b) [48].

#### 4.8.1. Pre-processing

The EEG data obtained from the 3 blocks of During were processed first. For every block, epochs were extracted from the EEG according to trial. Since the time to trigger electrical stimulation in the retention interval was different across trials, the length of epochs for different trials were also different. Then we deleted the epochs corresponding to the trials with wrong responses in the Sternberg task, and only the epochs for correct trials remained. Next, the detrending was performed to remove DC offsets and slow drifts (<1 Hz). Eye blink contaminations were then eliminated using an independent component analysis approach (implemented in EEGLAB). Epochs with residual eye movements or other artifacts were removed through visual inspection of the data. Because the trial numbers and the duration of trials obtained from the three stimulation conditions were quite different, it was necessary to balance the trial numbers and trial lengths of the three stimulation conditions. For each participant and each block, we deleted the shortest trials in the two conditions with more trials so that the trial numbers were the same as the condition with the minimum number of trials. Next, the epochs in each condition were sorted by the length of epochs from shortest to longest. Then we made the epochs with the same order among the three stimulation conditions have the same length by discarding the tail EEG data of the two longer epochs.

For pre-test and post-test EEG data, we matched the data lengths of the pre-test and post-test epochs with that of the epochs at During. For every participant and every condition, since the duration of all pre-test and post-test epochs was 2.5 s, we randomly assigned the length of all epochs from During to the epochs of pre-test and post-test, and deleted the tail EEG data of each epoch at pre-test and post-test based on the length assigned. Then the preprocessing of the EEG data from pre-test and post-test was the same as During.

#### 4.8.2. Analysis of alpha power (8-13 Hz)

For each epoch, the absolute spectrum was calculated using the Matlab function *pwelch*. Because of the different duration of different epochs, each epoch was first divided into several small segments with a duration of 500 ms. And there was an overlap of 400 ms from segment to segment. A fast Fourier transform (FFT) was calculated for each segment using a Hamming window and zero-padding to 2.048s. Then the spectra of the segments for every epoch were averaged to obtain the spectrum of the epoch. The resulting spectra of each block were averaged across epochs as well as across the alpha-bands (8-13Hz) per tACS-condition as the absolute power in alpha band. Finally, for subsequent statistical analysis, the relative alpha power of each block was calculated as the absolute power in alpha band divided by the absolute power across the frequency band of 1-45 Hz.

#### 4.8.3. Phase synchronization analysis

The phase lag index (PLI) is defined as

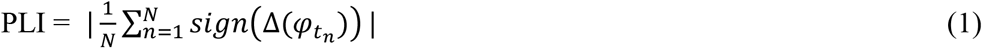

Where sign is the signum function that discards phase difference of 0 mod π, and Δ(*φ_t_n__*), *n* = 1 … *N* represents a time series of phase differences. PLI can obtain reliable estimates of phase synchronization against the presence of volume conduction [49]. Therefore, phase synchronization between Pz electrode and other 12 electrodes was estimated using the PLI method.

To obtain phase information, preprocessed data were bandpass-filtered in alpha-bands (8-13Hz) and Hilbert-transformed. The instantaneous phases could be extracted from the resulting complex values. The phase lag index (PLI) between Pz electrode and one of the other 12 electrodes was then computed for every epoch. Within every block, PLIs were then averaged across epochs in each tACS-condition. Finally, the average PLI between the alpha activity of the 7 frontal electrodes (Fp1, Fp2, F7, F3, Fz, F4, F8) and Pz electrode was used as an indicator of frontoparietal alpha synchronization.

### 4.9. Statistical analysis

The rate correct score (RCS) is defined as

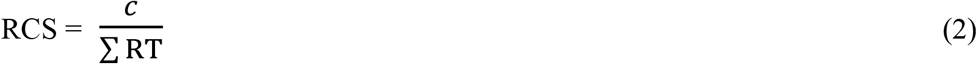

Where c is the number of correct responses, and the denominator refers to the sum of all reaction times (RTs) (including both correct and incorrect RTs) in the set of trials under consideration.

RCS was proposed in a memory task to integrate measurements of RT and accuracy into a single measure to incorporate meaning variance of both reaction time and accuracy [50]. It can be interpreted directly as the number of correct responses per unit time. This measure is more sensitive and less variable than single measures of RT and accuracy, especially under conditions in which an external manipulation or intervention causes parallel changes in reaction time and accuracy (i.e., both disturb or both improve) [51]. Follow-up researches further support the utility of RCS in cognitive science [52, 53].

Due to the suggested different mechanisms underlying online tACS effects [18, 20] (effects observed during stimulation) and offline tACS effects [26] (aftereffects beyond stimulation), we analyzed the online tACS effects firstly by comparing the online behavioral and EEG effects induced by in-phase and anti-phase tACS, then the offline tACS effects. To exclude the influences of pre-test, we normalized all metrics by subtracting the corresponding pre-test values.

The statistical analysis was performed using MATLAB 2016a. For statistical significance analysis, we used nonparametric permutation test throughout the manuscript to avoid the assumption of normality. For comparisons between the two paired groups, we used the permuted paired *t*-test statistic wherein we randomly mixed values from the two groups 5000 times to create a distribution of paired *t* values and computed an empirical p value from this distribution. Note that *t*-test statistics and degree of freedom are provided for reference only, and the reported p-values may differ from those expected from the *t*-distribution. The effect size (Cohen’s d-value) was calculated via G*Power 3.1 software [45]. Correlation analysis between two variables was calculated using permutation test based on Pearson’s linear correlation coefficient (r-value, two-tailed). We used the permuted Pearson’s correlation coefficients wherein we hold one variable constant and randomly permuted the other variable 5000 times to create a distribution of r values and computed an empirical p value from this distribution. To test for differences between two correlations, the obtained correlation coefficients were converted into z-values with Fisher’s r-to-z transformation so that the z scores can be analyzed for statistical significance.

### Data and code availability

The original data for analysis and custom Matlab code are available from the corresponding author (X.Z.) upon reasonable request.

## Acknowledgments

We thank Lizhuang Yang, Guanbao Cui, Xianze Zheng, Zhen Miao, Kaining Zhang and Zehua Fang for their assistance with experimental design. This work was supported by grants from The National Key Basic Research Program (2018YFC0831101), The National Natural Science Foundation of China (31771221, 71942003, 61773360, 31800927, 31900766 and 71874170), Major Project of Philosophy and Social Science Research, Ministry of Education of China (19JZD010), CAS-VPST Silk Road Science Fund 2021 (GLHZ202128), Collaborative Innovation Program of Hefei Science Center, CAS (2020HSC-CIP001). A portion of the numerical calculations in this study were performed with the supercomputing system at the Supercomputing Centre of USTC.

## Author contributions

X.Z. conceived the experiments; X.C., R.M., Q.W., and A.Y. performed the experiments; X.C., R.M., and W.Z. analyzed the data; R.M., X.X., and J.C. designed and realized the online phase-corrected closed-loop tACS system; X.C., R.M., and X.Z. wrote the manuscript; A.T.S., M.M.L., N.X.Y, S.W., Z.D., G.Q.Z., Q.C., and J.B. helped to polish the writing.

## Materials & Correspondence

Correspondence and requests for materials should be addressed to X.Z.

## Abbreviations

IAF: Individual Alpha Frequency
RCS: Rate Correct Score
tACS: Transcranial Alternating Current Stimulation
WM: Working Memory

