## Supplemental material for "Alpha Oscillatory Activity Causally Linked to Working Memory Retention: Insights from Online Phase-corrected Closed-loop Transcranial Alternating Current Stimulation (tACS)"

**Contents**

1. **More details for materials and methods**
   1.1 **The details of online phase-****corrected closed-loop tACS**
   1.2 **IAF and threshold determination**
   1.3 **Analysis of the phase alignment between EEG signal and tACS waveform**
   1.4 **Analysis of tACS questionnaire**
2. **The online comparison of accuracy and RT between in-phase tACS and anti-phase tACS**
3. **Supplementary power analysis**
   3.1 **The comparison of power in full frequency band between in-phase alpha-**

**tACS and anti-phase alpha-tACS**

3.2 **The comparison of individual alpha power (IAF ± 2Hz) between in-phase**

**tACS and anti-phase tACS**

4. **The online comparison of WPLI between in-phase tACS and anti-phase tACS**
5. **Complementary analysis of the behavior and EEG data in random-phase tACS**
   5.1 **The behavioral effects of random-phase tACS compared with in-phase**

**and anti-phase tACS**

5.2 **The EEG results of random-phase tACS compared with in-phase and**

**anti-phase tACS**

5.3 **The correlations between random-phase tACS induced changes in behavioral performance and alpha activities**

6. **The offline comparison between in-phase tACS effects and anti-phase tACS effects**
7. **In-phase and anti-phase tACS at the theta frequency**
8. **Supplementary discussion**
   8.1 **A discussion on the position of the stimulation electrode**
   8.2 **A discussion on the non-significance of in-phase tACS-induced**

**enhancement effects**

**1. More details for materials and methods**

**1.1. The details of online phase-corrected closed-loop tACS**

Illustrated in **Figure 1**, the online phase-corrected closed-loop tACS system included an EEG instrument, computation module and a custom designed tACS stimulator which could receive the instructions from the computation module. The closed-loop brain stimulation algorithm for online phase-corrected closed-loop tACS was introduced that could record brain oscillations by an EEG instrument, analyze the raw data online by computational algorithms to extract the phase of underlying brain rhythms, deliver tACS with a desired phase difference from the endogenous targeted brain oscillations based on the computational outcome, and closing the loop, affect the neuronal activity.

1.1.1. ***Signal processing***

The EEG signal of interested electrode (Pz electrode for our study) was stored in a moving 500 ms time window. Because of the performance setting of EEG amplifier, the EEG data was updated every 10ms, and the analysis should be finished within this time range. First, the recorded EEG signal over the time window was bandpass filtered to extract the activity within the frequency range of interest (IAF ± 2Hz for alpha-tACS). A 200th order finite impulse response (FIR) filter was used because of its linear-phase property, and the raw data was reversed before filter to maintain the latest data after phase correction. Then, whether there were peaks or troughs in the last 10ms of the filtered signal were estimated based on the fact that the peaks and troughs are local maximums or minimums.

In in-phase tACS, stimulation was triggered when two consecutive peaks exceeded the threshold (in our study, threshold was calculated previously using the EEG signal of the pre-test block, see Experimental procedures section); while in the condition of anti-phase tACS, the stimulation was triggered when the absolute value of two consecutive troughs exceeded the threshold. In theory, in both in-phase and anti-phase tACS, the algorithm waited for a time interval of 3/4 cycles at the frequency of interest (IAF for our study) when the stimulation trigger requirements were met, and then the stimulation was applied to the participants by tACS stimulator. To correct for the time delays in the system introduced by the signal processing or hardware delays, the delay time was experimentally measured and corrected (see more details in Supplementary material: 1.1.2. Delay correction section). The onset phase of the sine-wave stimulation was invariably 0 in all conditions. As a result, the phase difference between in-phase tACS and the endogenous alpha activity was 0°, while the phase difference between anti-phase tACS and the endogenous alpha activity was 180°.

In random-phase tACS, to make the phase difference not fixed across trials, we artificially set whether to apply tACS during the retention interval of a certain trial and the start time of tACS within each trial. 5 participants were recruited to complete a pretest in which every participant performed two stimulation sessions: in-phase tACS and anti-phase tACS. Their stimulation sequences (whether to apply tACS in each trial) and the start times of tACS within each trial (the start time of tACS relative to the onset of the retention interval if stimulation was triggered in a trial) were collected. When a participant performed random-phase session in the formal experiment, we randomly assigned one of the 10 stimulation sequences to the participant. Then if tACS should be applied in a trial, we specified the start time of tACS within the WM retention interval of the trial by randomly selecting one of the start times of tACS collected in the pretest.

1.1.2. ***Delay correction***

In in-phase and anti-phase tACS, when the alpha activity reached the threshold for triggering tACS, the application time of tACS stimulation should be precisely controlled so that the phase difference between the tACS waveform and the endogenous alpha oscillations was 0° or 180°. However, it should be noted that signal processing components, such as the hardware filters used in EEG recording instruments, and the software filters applied to the signal during processing, could introduce frequency or cycle time dependent delays to the recordings. Additionally, hardware delays in communicating the stimulation instructions to the stimulator were also inevitable.

We addressed this issue by experimentally measuring the time delays introduced by all the components of the system, and then applying a time correction to achieve the desired phase difference between tACS waveform and the endogenous brain oscillations. To obtain and calibrate the delays, we used a 50Ω resistor to simulate the resistance of human brain and a signal generator (XD2, Zhonghuan Keyi Electronic Instrument Co., Ltd, Tianjin, China) to simulate the EEG signal generated by the brain. We then started the online phase-corrected closed-loop tACS system, connected the output of the signal generator and the output of the tACS stimulator to the EEG recording instrument, and then analyzed the collected data to obtain the delays of the tACS waveform relative to the simulated EEG signal.

We randomly selected 10 frequencies that may be used in the formal experiments within the frequency range of 8-13 Hz and tested the time delays in both in-phase and anti-phase tACS conditions. For each frequency, we applied online phase-corrected closed-loop tACS stimulation at that frequency with a duration of 0.5s for 100 times, and recorded the tACS waveform and the sinusoidal waveform at the same frequency generated by the signal generator continuously. The voltage of tACS was about 8.75mV, lower than the saturation voltage of the EEG amplifier. We then computed the delays between the first peak of each tACS waveform segment and the nearest peak of signals generated by the signal generator in in-phase tACS, or computed the delays between the first peak of each tACS waveform segment and the nearest through of the simulated EEG signals in anti-phase tACS, which was actually the system delays.

At the same frequency, there was no difference between the system delays measured in in-phase tACS and anti-phase tACS. Furthermore, we found that the delays were positively correlated with the cycle time of the tACS waveform (r = 0.999, p <0.001) (**Figure S1**). The delays could be estimated from the formula: L = 1.2013*(1000/f)-34.8, where L was the delays and f was the tACS frequency. The unit of L was milliseconds, and 1000/f represented the cycle time of the tACS waveform at the frequency of f Hz. In the formal experiment, we used the formula above to estimate the system delays for a given tACS frequency and subtracted the delays from the waiting time when tACS was triggered.

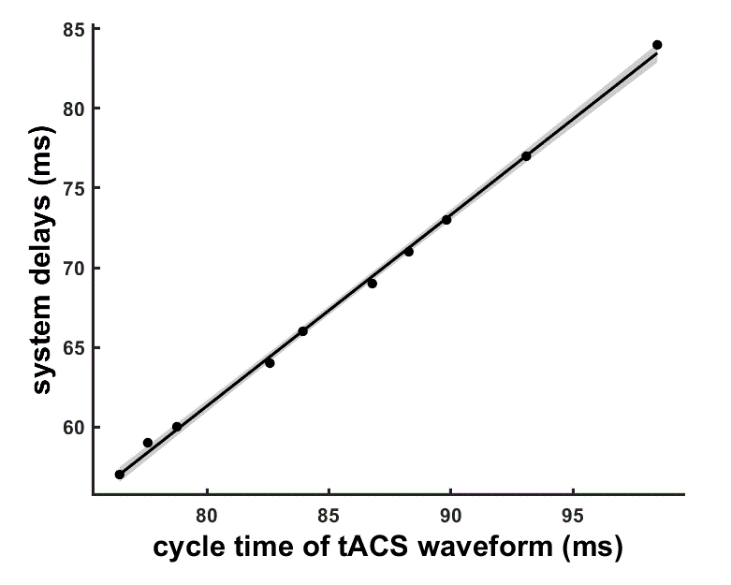

**Figure S1. The correlation between the system delays of the tACS system and the cycle time of tACS waveform.** The system delays were positively correlated with the cycle time of the tACS waveform.

1.1.3. ***Design of tACS stimulator***

To apply tACS dependent on the brain states, we designed a tACS stimulator which could communicate with the computer through serial USB. To protect participants from any potential harm, voltages of the output port were monitored by protect circuits to ensure security. Once the voltage exceeded the safety threshold (36 V, peak to peak), the output port would be cut down by the relay. The tACS stimulator is powered by four rechargeable batteries and isolated from mains electricity. The computer communicating with the tACS stimulator also stopped charging once the phase-corrected closed-loop tACS was connected with participants.

Illustrated in **Figure 1B**, the main part of the stimulator consisted of an Arduino Uno microcontroller board, a digital-to-analog converter and two operational amplifiers. The Arduino Uno microcontroller board was the core control part of our designed stimulator. The timing of the next tACS calculated by the computer was communicated to an Arduino Uno Microcontroller board (Arduino) through serial USB communication. Then, the stimulator follows the Arduino-generated voltage waveforms and produces a current controlled output proportional to that voltage.

The specifications of the custom designed tACS stimulator were illustrated in **table S1**.

**Table S1. The specifications of the custom designed tACS stimulator**

| Number of channels | 1 pair (1 anode, 1 cathode) |
| --- | --- |
| Power Supply | rechargeable battery |
| Maximum Duration | 3 hours |
| Maximum Voltage | ±18V |
| Stimulation Current (peak-to-peak) | max. 4mA (increment: 0.02mA) |
| Frequency | 0.5-100Hz (increment: 0.01Hz) |

1.2. **IAF and threshold determination**

To determine the IAF and the threshold for triggering tACS, the EEG data at pre-test was analyzed. The first 0.5s EEG data in each trial at pre-test was discarded due to the weak alpha oscillations, and the remaining EEG data at Pz electrode was analyzed. To this end, there were sixty 2s segments. Subsequently, the segments corresponding to the error trials were deleted. Preprocessing included the following steps: detrending, blink artifacts correction (using independent component analysis approach), rejecting all segments containing activity exceeding a threshold of 100 uV, and the removal of segments with residual eye blinks or other artifacts. After preprocessing, a fast Fourier transformation (FFT, frequency resolution 0.1953 Hz) was performed for each remaining segment, and the resulting spectra were averaged. The prominent alpha peak (8-13 Hz) was detected and its frequency was defined as IAF.

To really target the interested endogenous oscillations (*i.e.*, alpha oscillations), we aimed to deliver tACS only when strong endogenous interested oscillations exist as previous closed-loop tACS studies did [1, 2]. Therefore, we calculated a threshold for triggering tACS. We performed an offline analysis of the pre-test EEG data recorded in the retention interval of each trial the same as the online EEG monitoring (see more details in Supplementary material: 1.1.1. Signal processing). The window length was 500ms and the sliding step was 10ms. Bandpass filtering was performed on the EEG signals in each time window to extract the activity within the frequency range of IAF ± 2Hz. Then we decided whether a peak or trough existed in the last 10 ms of the alpha oscillations extracted from each time window, and recorded the detected peak or trough values. The lower quartile of all peak values and the lower quartile of all absolute values of troughs were used as the threshold for in-phase tACS and anti-phase tACS, respectively.

1.3. **Analysis of the phase alignment between EEG signal and tACS waveform**

Because of the unstable phases of EEG signals, we analyzed the phase alignment between EEG signals at pre-test and tACS waveform generated offline using the method same as online stimulation to test the performance of our system. The EEG signal was bandpass filtered within the frequency range of IAF±2Hz by 200th order FIR filter. The phases of filtered EEG (φEEG) and tACS waveform (φtACS) were calculated by Hilbert transformation and compared using the phase locking value (PLV) and phase differences: PLV = abs(mean(exp(i*(φEEG-φtACS)))); phase differences = angle(mean(exp(i*(φEEG-φtACS)))). The item exp(i*(φEEG-φtACS)) was averaged across time points in a single trial and then averaged across trials in a single experiment session.

PLV of in-phase and anti-phase conditions were all significant when compared to 0 (permuted one sample *t*-test, in-phase: t(38) = 11.607, p < 0.001; anti-phase: t(38) = 11.322, p < 0.001), while the difference between in-phase and anti-phase condition in PLV was not significant (permuted paired *t*-test, t(38) = 0.559, p = 0.581). Analysis of the phase differences was performed using the circular statistics toolbox [3]. To test if the phase differences would change over time, we sort the time points into 6 groups, which were 1-100ms, 101-200ms, 201-300ms, 301-400ms, 401-500ms and 501ms to the end of the tACS waveform. Harrison-Kanji test indicated that the main effect of tACS condition was significant (p<0.001) while the main effect of time was not significant (p = 0.411). The above results indicated that despite of the unstable phases of EEG signals, the phase alignment between EEG signals and tACS waveform within 0.8 s is as expected.

1.4. **Analysis of tACS questionnaire**

After tACS finished, participants completed a questionnaire to assess ten possible side-effects of the tACS stimulation by rating from 0 (none) to 4 (strong) the intensity of: itching, pain, burning, warmth/heat, pinching, metallic/iron taste, fatigue, dizziness, nausea, phosphenes, or any other side-effects perceived. If a certain side-effect was present, participants were asked to evaluate the possibility that the side-effect was related to tACS by rating from 0 (none) to 4 (definite).

In order to provide an evaluation of the general perceived discomfort induced by tACS, two new aggregate variables were proposed: discomfort_1 and discomfort_2. Discomfort_1 was computed as the summation of the intensity score recorded for each single sensation, so that the discomfort_1 variable ranged from 0 (absence of discomfort) to 40 (maximum discomfort). Discomfort_2 was defined as the summation of the intensity score for each single sensation multiplied by the possibility score of the sensation related to tACS, ranging from 0 to 160.

There were no differences in the general discomfort between the tACS conditions. Permuted paired *t*-tests showed no difference between any two conditions for either discomfort_1 (in-phase vs anti-phase: t(38) = 0.956, p = 0.370; in-phase vs random-phase: t(38) = -0.259, p = 0.820; anti-phase vs random-phase: t(38) = -1.212, p = 0.249) or discomfort_2 (in-phase vs anti-phase: t(38) = 0.044, p = 0.971; in-phase vs random-phase: t(38) = -0.481, p = 0.646; anti-phase vs random-phase: t(38) = -0.611, p = 0.563). These results demonstrated that participants were not able to distinguish the three tACS conditions and excluded the possibility that the difference between the effects of in-phase and anti-phase tACS was attributed to the difference of tACS-induced discomfort.

2. **The online comparison of accuracy and RT between in-phase tACS and anti-phase tACS**

We used paired *t*-tests to explore whether anti-phase tACS induced lower accuracy and slower response time compared to in-phase tACS during stimulation, as we hypothesized that anti-phase tACS impaired WM performance compared to in-phase tACS. Compared to in-phase tACS, anti-phase tACS induced a marginally significant reduction in accuracy (permuted paired *t*-test, t(38) = -1.958, p = 0.057, Cohen’s d = 0.314) (**Figure S2A**), which further supports the different modulation effects of in-phase tACS and anti-phase tACS on WM performance. No significant difference in RT (permuted paired *t*-test, t(38) = 0.21, p = 0.840) was found between in-phase tACS and anti-phase tACS (**Figure S2B**).

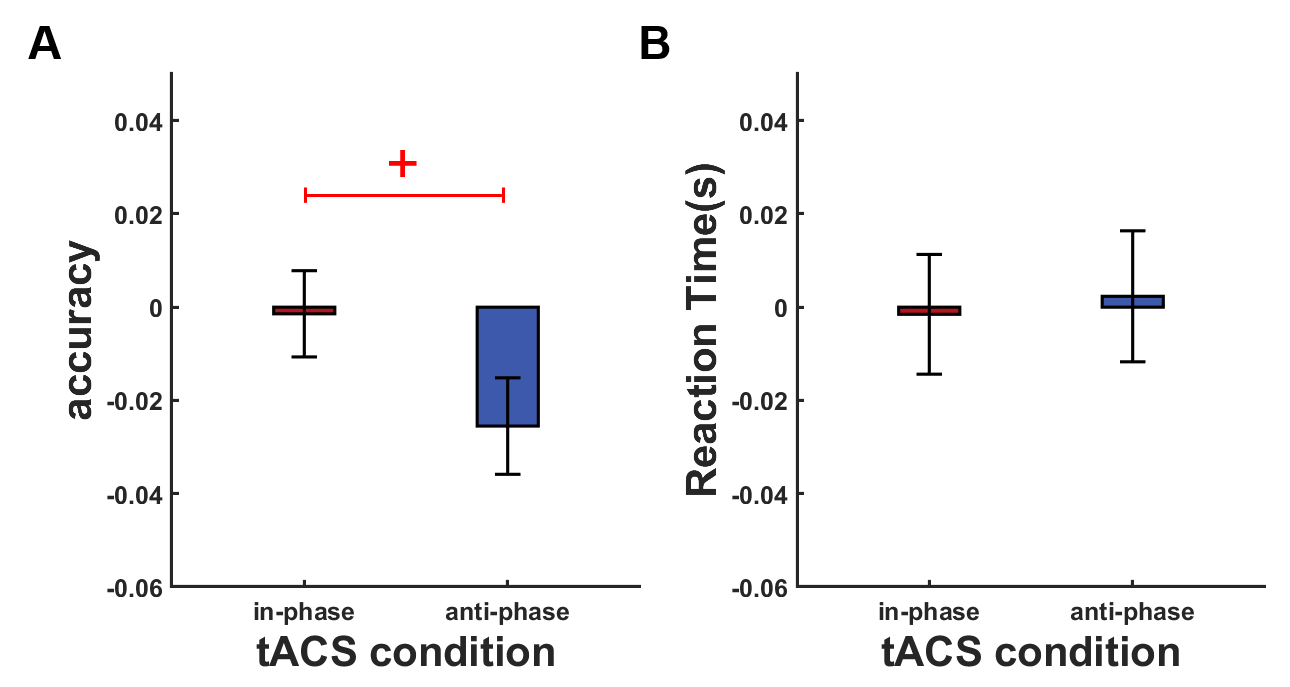

**Figure S2. The online effects of tACS on** (**A**) accuracy, and (**B**) reaction time at During for the two stimulation conditions: in-phase and anti-phase tACS. Accuracy and reaction time are given relative to pre-test. Error bars represent SEM; + marginally significant at 0.05< p <0.1.

3. **Supplementary EEG power analysis**
   3.1. **The comparison of power in full frequency band between in-phase alpha-tACS and anti-phase alpha-tACS**

Many previous studies have shown the frequency-specific modulation of tACS in both animals [4] and humans [5]. To further support the modulation effects of our online phase-corrected closed-loop tACS on brain activity, we tested the influences of this innovative tACS system on the full physiological frequency band of the EEG (from 1 Hz to 45 Hz). By convention, the waking EEG was divided into five distinct frequency bands: delta (1-4Hz), theta (4-8Hz), alpha (8-13Hz), beta (13-30Hz) and gamma (30-45Hz). We computed the relative power of each frequency band, with the analysis method same as alpha power (See detailed information in Methods). Permuted paired *t*-tests were performed to compare the differences between in-phase and anti-phase tACS within each frequency band. No significant differences were found in any of the frequency bands except for the alpha band at During (permuted paired *t*-test, delta: t(38) = 1.204, p = 0.224; theta: t(38) = 0.154, p = 0.881; alpha: t(38) = -2.257, p = 0.027; beta: t(38) = 0.124, p = 0.904; gamma: t(38) = 0.994, p = 0.343) (**Figure S3**). As hypothesized, these results demonstrated that the modulation of the endogenous brain oscillations was restricted to the tACS frequency.

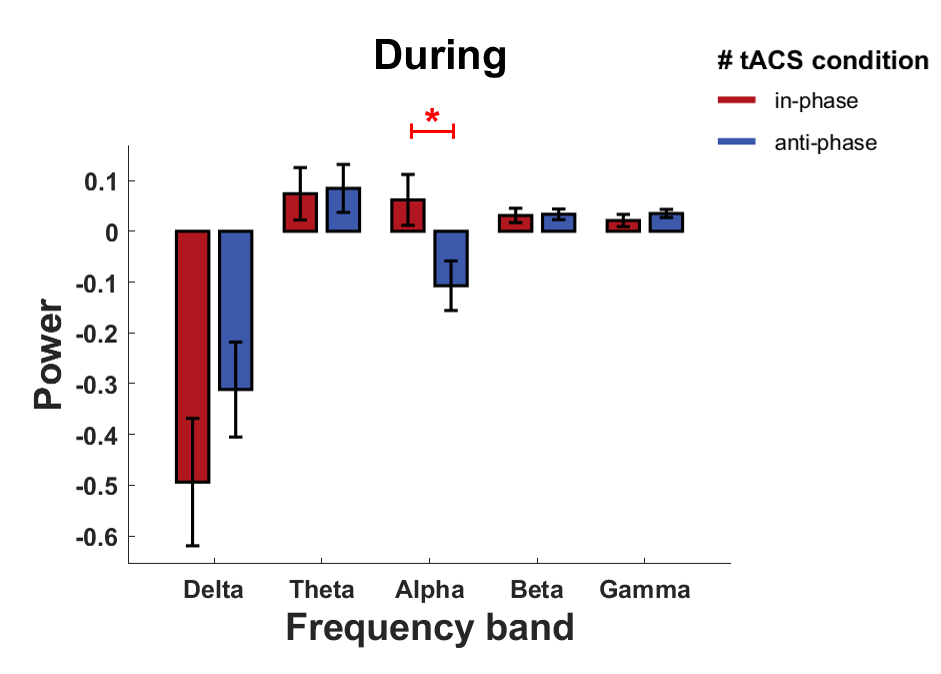

**Figure S3. The comparison of power in delta, theta, alpha, beta and gamma bands between in-phase and anti-phase alpha-tACS at During**. Powers are given relative to the corresponding values in pre-test. Error bars represent the SEM; *significant at p <0.05, ** significant at p <0.01.

3.2. **The comparison of individual alpha power (****IAF ± 2Hz) between in-phase tACS and anti-phase tACS**

Considering that the frequency of our alpha-tACS was calculated based on individual traits (IAF), we also analyzed the impact of alpha tACS on the individual alpha power within the frequency range of IAF ± 2Hz at Pz electrode given that alpha oscillations within this frequency band were monitored during closed-loop tACS, with the analysis method same as alpha power (8-13 Hz) (See detailed information in Methods). Permuted paired t-tests were performed to compare the differences in individual alpha power between in-phase and anti-phase tACS. Anti-phase tACS marginally significantly suppressed the alpha power at Pz electrode compared to in-phase tACS at During (permuted paired *t*-test, t(38) = -1.811, p = 0.077, Cohen’s d = 0.290) (**Figure S4**). This result is consistent with the result about alpha power (8-13Hz), further supporting the modulation effects of alpha-tACS on alpha oscillations.

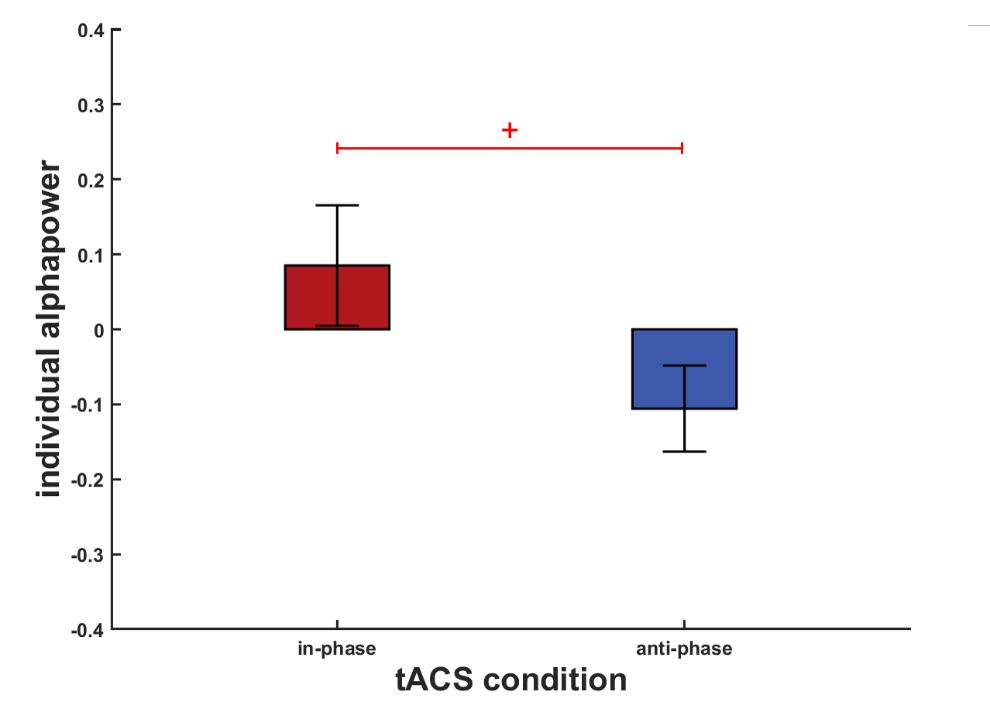

**Figure S4. The online effects of alpha-tACS on individual alpha power (IAF ± 2Hz) at During for the two stimulation conditions: in-phase and anti-phase tACS.** Individual alpha power is given relative to pre-test. Error bars represent SEM; + marginally significant at 0.05< p <0.1.

4. **The online comparison of WPLI between in-phase tACS and anti-phase tACS**

To further illustrate the reliability of our results, we chose another index from the family of phase-synchronization indices, weighted phase lag index (WPLI) [6], to measure frontoparietal alpha synchronization. WPLI may be more sensitive to additional, unrelated noise resources. We found that compared to in-phase tACS, anti-phase tACS marginally significantly disturbed frontoparietal alpha synchronization at During (permuted paired *t*-test, t(38) = -1.803, p = 0.075, Cohen’s d = 0.289) (**Figure S5**). This result is consistent with the result calculated by PLI, suggesting that our online phase-corrected closed-loop tACS modulate the connectivity of distributed brain regions.

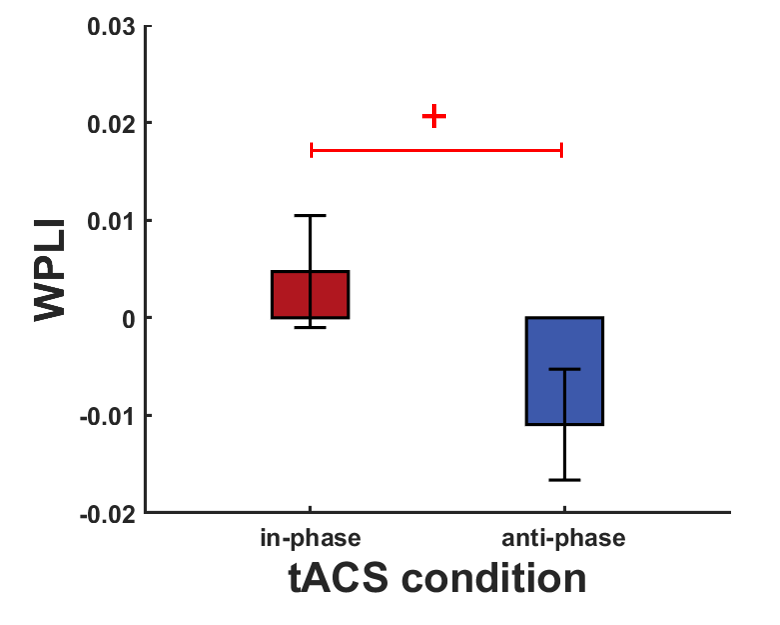

**Figure S5. The online effects of tACS on frontoparietal alpha synchronization, indexed by WPLI at During for the two stimulation conditions: in-phase and anti-phase tACS.** WPLI is given relative to pre-test. Error bars represent SEM. + marginally significant at 0.05< p <0.1.

5. **Complementary analysis of the behavior and EEG data in random-phase tACS**
   5.1. **The behavioral effects of random-phase tACS compared with in-phase and anti-phase tACS**

To explore the effects of the control condition, here we compared the effects of random-phase tACS with both in-phase and anti-phase tACS using permuted paired *t*-tests. At During, compared with random-phase tACS, either in-phase or anti-phase tACS didn’t show significant differences in RCS (permuted paired *t*-test, in-phase vs random-phase: t(38) = 1.464, p = 0.146; anti-phase vs random-phase: t(38) = -0.444, p = 0.661), accuracy (permuted paired *t*-test, in-phase vs random-phase: t(38) = 1.043, p = 0.302; anti-phase vs random-phase: t(38) = -0.659, p = 0.521) and RT (permuted paired *t*-test, in-phase vs random-phase: t(38) = -0.186, p = 0.853; anti-phase vs random-phase: t(38) = 0.036, p = 0.975) (**Figure S6**). Despite the lack of significant differences, the three behavioral metrics of random-phase tACS were all between that of in-phase tACS and anti-phase tACS at the stimulation period.

The behavioral difference between in-phase tACS, anti-phase tACS and random-phase tACS also didn’t reach significance at post-test (permuted paired *t*-test, RCS: in-phase vs random-phase: t(38) = -0.153, p = 0.877; anti-phase vs random-phase: t(38) = -0.017, p = 0.986. accuracy: in-phase vs random-phase: t(38) = -0.608, p = 0.540; anti-phase vs random-phase: t(38) = 0.278, p = 0.782. RT: in-phase vs random-phase: t(38) = 0.215, p = 0.838; anti-phase vs random-phase: t(38) = 0.348, p = 0.722) (**Figure S6**).

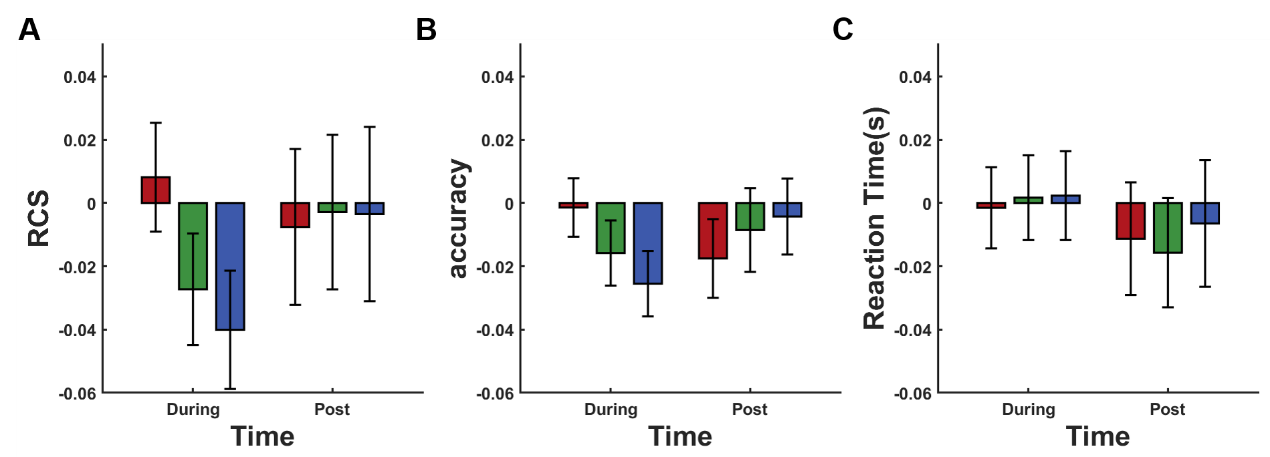

**Figure S6. The effects of tACS on working memory performance.** The change for (**A**) rate correct score (RCS), (**B**) accuracy, and (**C**) reaction time at During and post-test for the three stimulation conditions: in-phase, random-phase and anti-phase tACS. RCS, accuracy and reaction time are given relative to pre-test (subtract corresponding pre-test values). Error bars represent SEM.

5.2. **The EEG results of** **random-phase tACS compared with in-phase and** **anti-phase tACS**

At During, the alpha power at Pz electrode in random-phase tACS was also between in-phase tACS and anti-phase tACS (permuted paired *t*-test, During: in-phase vs random-phase: t(38) = 0.231, p = 0.819; anti-phase vs random-phase: t(38) = -2.631, p = 0.013). At post, paired *t*-tests demonstrated a significant increase of alpha power in in-phase tACS compared with random-phase tACS (permuted paired *t*-test, in-phase vs random-phase: t(38) = 2.585, p = 0.008), but no significant difference was observed in anti-phase tACS (permuted paired *t*-test, anti-phase vs random-phase: t(38) = 1.314, p = 0.193) (**Figure S7A**).

The change of frontoparietal alpha synchronization induced by random-phase tACS was also between in-phase tACS and anti-phase tACS during tACS and at post-test (permuted paired *t*-test, During: in-phase vs random-phase: t(38) = 0.747, p = 0.456; anti-phase vs random-phase: t(38) = -1.824, p = 0.079. post: in-phase vs random-phase: t(38) = 0.961, p = 0.341; anti-phase vs random-phase: t(38) = -0.124, p = 0.906) (**Figure S7B**). This indicated that the physiological effects of random-phase tACS are also between in-phase and anti-phase tACS as hypothesized despite of the small effect size.

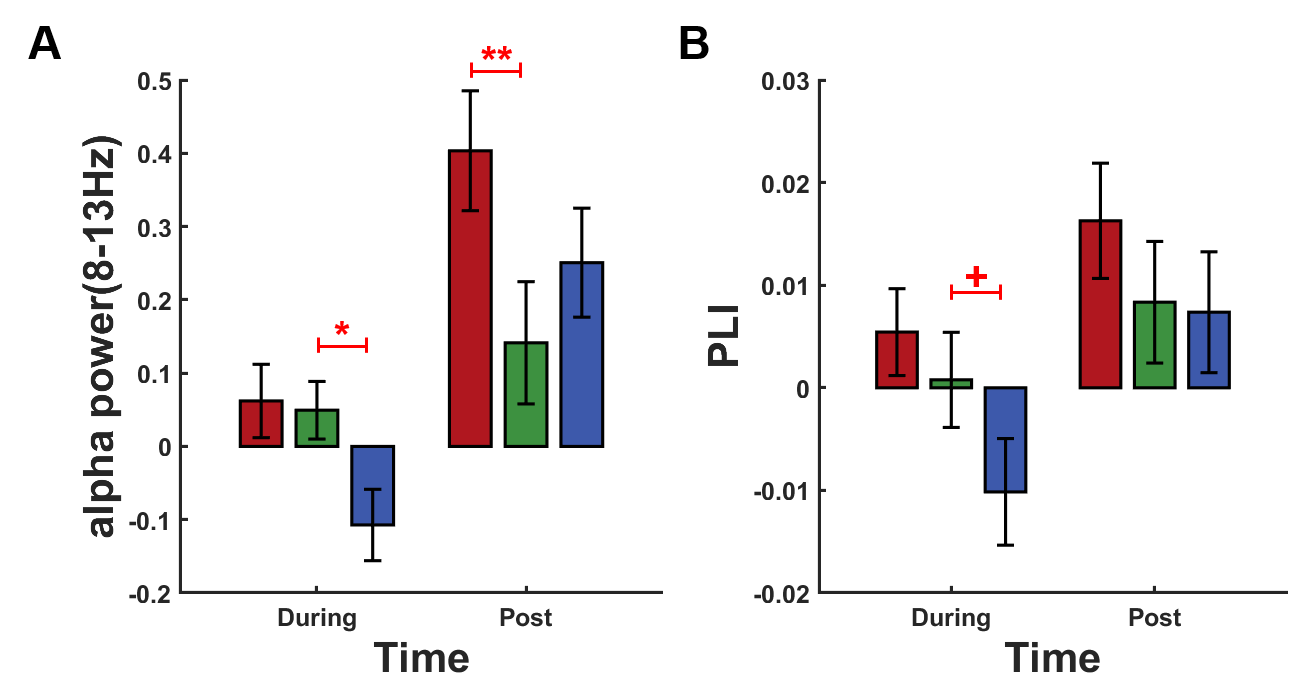

**Figure S7. EEG results elicited by in-phase, random-phase and anti-phase tACS conditions.** (**A**) the alpha power of Pz electrode in in-phase and anti-phase tACS compared with random-phase tACS. (**B**) the frontoparietal alpha synchronization, indexed by PLI in in-phase and anti-phase tACS compared with random-phase tACS. Alpha power and frontoparietal alpha synchronization are given relative to pre-test (subtract corresponding pre-test values). Error bars represent SEM; + marginally significant at 0.05< p <0.1, *significant at p <0.05, ** significant at p <0.01.

5.3. **The correlations between random-phase tACS induced changes in behavioral performance and alpha activities**

For random-phase tACS, changes in alpha power and RCS were not related during and after tACS (permuted Pearson’s correlation: During: r = -0.066, p = 0.695; post: r = -0.074, p = 0.653). What’s more, the correlation coefficients of random-phase tACS were between that of in-phase tACS and anti-phase tACS, which supports that the positive correlation between alpha power and WM performance was not merely due to the general effects of tACS (**Figure S8A**).

No correlation between changes in frontoparietal alpha synchronization and changes in RCS was observed for random-phase tACS during and after tACS (permuted Pearson’s correlation, During: r = 0.021, p = 0.935; post: r = -0.165, p = 0.309). In consistent with the correlation between changes in alpha power and changes in RCS, the correlation coefficient of random-phase tACS was also between that of in-phase tACS and anti-phase tACS, which supports that not tACS but in-phase tACS induced the increase of frontoparietal alpha synchronization and further helped to improve WM performance (**Figure S8B**).

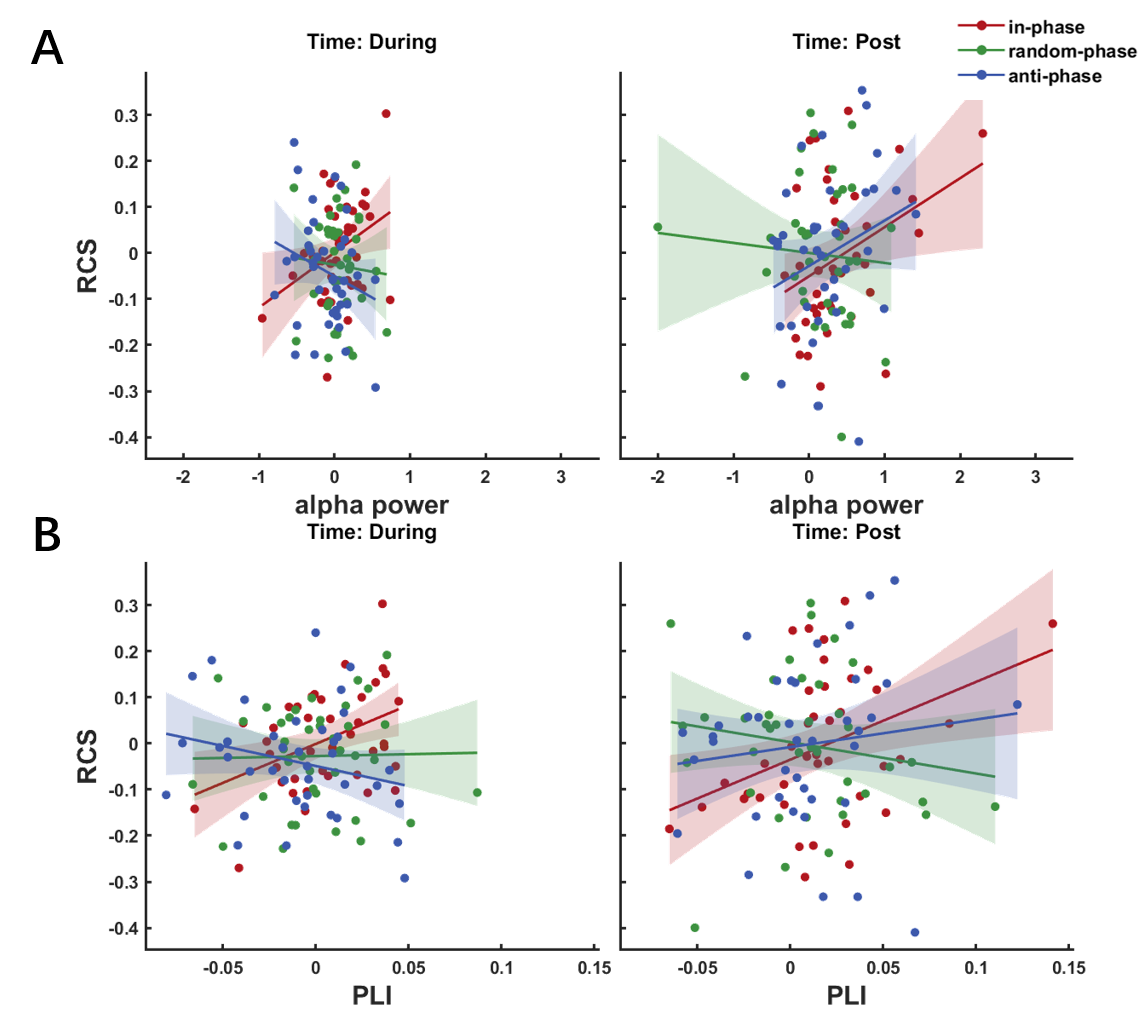

**Figure S8. Relationship between tACS-induced changes in RCS and EEG metrics for in-phase, random-phase and anti-phase tACS.** (**A**) Correlation between the changes in RCS and parietal alpha power relative to pre-test at each time point (During and post). (**B**) Correlation between the changes in RCS and frontoparietal alpha synchronization at each time point (During and post). RCS, alpha power and frontoparietal alpha synchronization are given relative to pre-test (subtract corresponding pre-test values).

6. **The offline comparison between in-phase tACS effects and anti-phase tACS effects.**

Finally, we explored whether anti-phase tACS induced offline suppression in WM performance and alpha activity compared to in-phase tACS using permuted paired *t*-tests. Alpha power in anti-phase tACS was marginally significantly weaker than in-phase tACS (permuted paired *t*-test, t(38) = -1.704, p = 0.094, Cohen’s d = 0.273) . But unfortunately, compared to in-phase tACS, anti-phase tACS didn’t suppress WM performance (permuted paired *t*-test, RCS: t(38) = 0.123, p = 0.900; accuracy: t(38) = 0.941, p = 0.325; RT: t(38) = 0.178, p = 0.860), or frontoparietal alpha synchronization (permuted paired *t*-test, t(38) = -1.105, p = 0.284) (**Figure S9**).

Then we further explored the changes from pre-test to post-test within in-phase and anti-phase tACS. Compared to pre-test, no suppression in anti-phase tACS was observed at post-test (permuted paired *t*-test, RCS: t(38) = -0.125, p = 0.898; accuracy: t(38) = -0.351, p = 0.747; RT: t(38) = -0.319, p = 0.754; alpha power: t(38) = 3.327, p = 0.002; PLI: t(38) = 1.235, p = 0.242). For in-phase tACS, there were also no significant increase in WM performance, but significant increase in alpha power and frontoparietal alpha synchronization at post-test (permuted paired *t*-test, RCS: t(38) = -0.303, p = 0.756; accuracy: t(38) = -1.392, p = 0.154; RT: t(38) = -0.626, p = 0.535; alpha power: t(38) = 4.869, p < 0.001, Cohen’s d = 0.780; PLI: t(38) = 2.86, p= 0.008, Cohen’s d = 0.458). Considering the enhancement of alpha activities in both in-phase and anti-phase tACS, the significant increase in alpha activities at post-test may be due to the increased efforts needed to suppress irrelevant information after a long time of task performance [7], thereby not contributing to the improvement of WM performance.

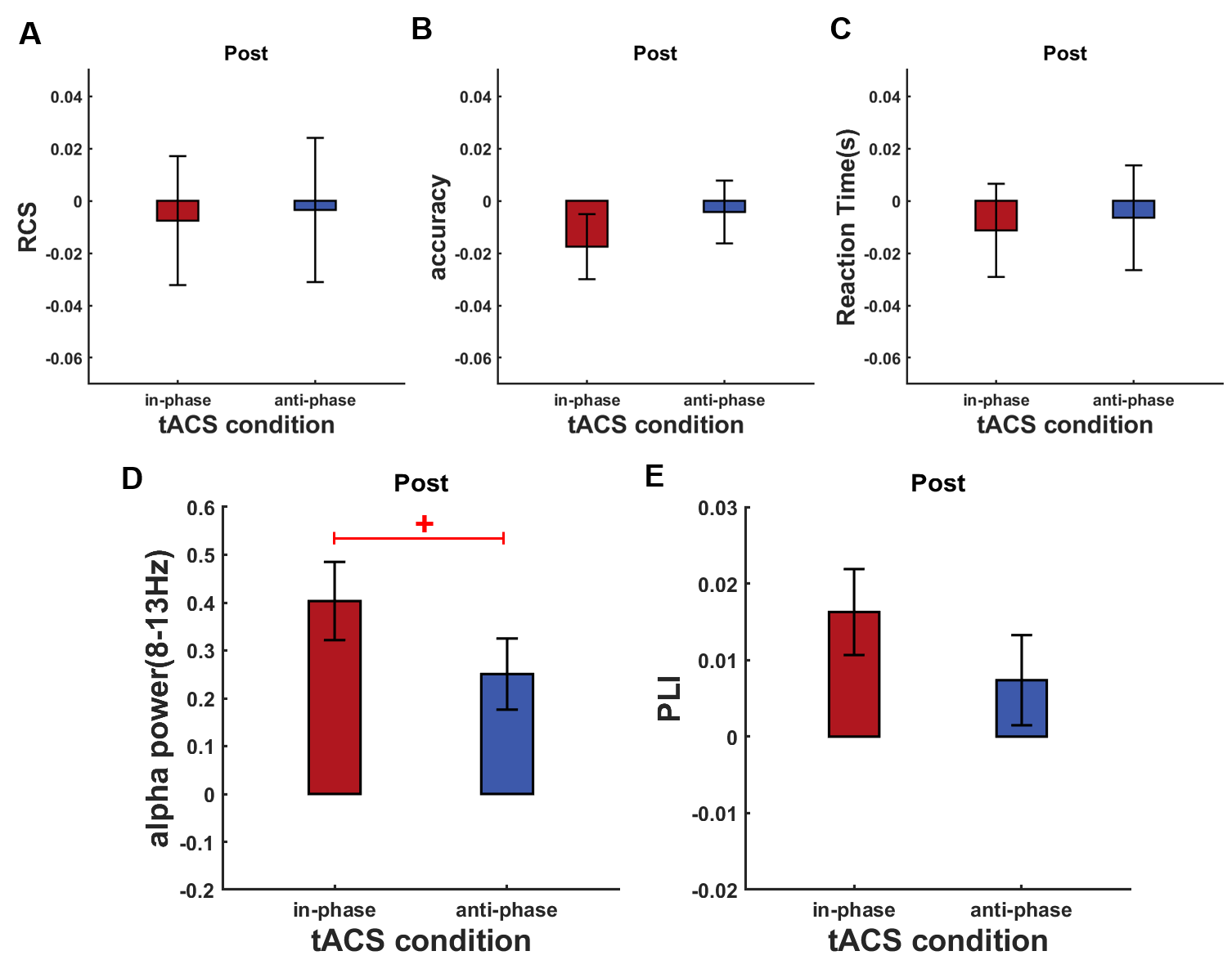

**Figure S9. The offline effects induced by in-phase tACS and anti-phase tACS in** (**A**) RCS, (**B**) accuracy, (**C**) RT, (**D**) alpha power and (**E**) frontoparietal alpha synchronization. RCS, accuracy, reaction time, alpha power and frontoparietal alpha synchronization are given relative to pre-test (subtract corresponding pre-test values). Error bars represent SEM; + marginally significant at 0.05< p <0.1.

7. **In-phase and anti-phase tACS at the theta frequency**
   7.1. **Participants**

39 volunteers participated in the theta tACS experiment. One participant was excluded from the analyses due to equipment failure to trigger intended electrical stimulation. The data of the remaining 38 participants were used for subsequent analysis (24 females, mean age ± SD：23.0 ± 1.9 years, mean education ± SD：16.8 ± 1.5 years). All participants gave written informed consent prior to the study.

7.2. **Experimental procedures and stimulation paradigm**

Each participant underwent 2 experimental sessions separated by at least 3 days at approximately the same time of the day. Using a single-blind within-subject design, participants received in-phase tACS or anti-phase tACS in each session, with the order of sessions counterbalanced across participants. The experimental procedure within each session was the same as alpha tACS, including the first pre-test block, During, and the final post-test block.

The stimulation configuration was also the same as alpha tACS except tACS frequency. After the first EEG block (pre-test), IAF was calculated using the EEG signal of pre-test. Then individual theta frequency (ITF) was then determined as (IAF – 5) Hz as some previous studies did [8]. At During, our online phase-corrected closed-loop tACS system was applied to deliver tACS at ITF when EEG signal met requirements.

The online phase-corrected closed-loop tACS system used in theta tACS was also same as alpha tACS except for two differences. First, the threshold was also calculated using the EEG signal of pre-test (see Supplementary material: 1.2. IAF and threshold determination described), but the 2.5% quantile rather than the lower quartile of all peaks or all absolute throughs was used as the threshold value. Due to the weak parietal theta oscillations and the fewer cycles of theta oscillations, we set lower threshold for theta tACS to make the stimulation numbers of theta tACS comparable to alpha tACS. Second, to minish the difference of the influence of band filter on the upper and lower bound of the target frequency band and make the phase determination of endogenous theta oscillations more accurate, the recorded EEG signal over the 500ms time window was bandpass filtered to extract the activity within 1 Hz (rather than 2 Hz) around ITF.

7.3. **Theta-tACS sensations and blinding**

Theta-tACS induced sensations were computed using the same method described in alpha-tACS (see Supplementary material: 1.4. Analysis of tACS questionnaire). There were no differences in the general discomfort between the tACS conditions. Permuted paired t-tests showed no difference between in-phase tACS and anti-phase tACS for either discomfort_1 (in-phase vs anti-phase: t(37) = -1.433, p = 0.182) or discomfort_2 (in-phase vs anti-phase: t(37) = -1.648, p = 0.117). This also demonstrated that participants were not able to distinguish in-phase tACS and anti-phase tACS in the control theta tACS experiment.

8. **Supplementary discussion**
   8.1. **A discussion on the position of the stimulation electrode**

The stimulation electrode in our study was not precisely located at the monitored Pz electrode, but between the Pz electrode and the Oz electrode to avoid the pollution of EEG signals. One may wonder whether tACS in our study was able to target the alpha activity recorded from Pz electrode. Alpha activity recorded from Pz electrode generated from the widely distributed sources in parieto-occipital regions [9], supporting the capacity of our tACS to target these sources.

8.2. **A discussion on the non-significance of in-phase tACS-induced enhancement effects**

The improvement effects on WM performance and alpha activity induced by in-phase tACS were not significant at During, so that one may question the enhancement effects of in-phase tACS. Please note that although not reaching the significance, in-phase tACS showed the tendency to enhance all of the three detected metrics (i.e., RCS, parietal alpha power and frontal-parietal alpha synchronization) during tACS. The lack of significant enhancement may be due to the fact that it is difficult for tACS to increase endogenous so strong alpha oscillations [10] induced by the difficult Sternberg paradigm.
